# *De-novo* assembly of zucchini genome reveals a whole genome duplication associated with the origin of the *Cucurbita* genus

**DOI:** 10.1101/147702

**Authors:** Javier Montero-Pau, José Blanca, Aureliano Bombarely, Peio Ziarsolo, Cristina Esteras, Carlos Martí-Gómez, María Ferriol, Pedro Gómez, Manuel Jamilena, Lukas Mueller, Belén Picó, Joaquín Cañizares

## Abstract

The *Cucurbita* genus (squashes, pumpkins, gourds) includes important domesticated species such as *C. pepo*, *C. maxima* and *C. moschata*. In this study, we present a high-quality draft of the zucchini (*C. pepo*) genome. The assembly has a size of 263 Mb, a scaffold N50 of 1.8 Mb, 34,240 gene models, includes 92% of the conserved BUSCO core gene set, and it is estimated to cover 93.0% of the genome. The genome is organized in 20 pseudomolecules, that represent 81.4% of the assembly, and it is integrated with a genetic map of 7,718 SNPs. Despite its small genome size three independent evidences support that the *C. pepo* genome is the result of a Whole Genome Duplication: the topology of the gene family phylogenies, the karyotype organization, and the distribution of 4DTv distances. Additionally, 40 transcriptomes of 12 species of the genus were assembled and analyzed together with all the other published genomes of the Cucurbitaceae family. The duplication was detected in all the *Cucurbita* species analyzed, including *C. maxima* and *C. moschata*, but not in the more distant cucurbits belonging to the *Cucumis* and *Citrullus* genera, and it is likely to have happened 30 ± 4 Mya in the ancestral species that gave rise to the genus.

## Introduction

*Cucurbita pepo* L. is the main crop of the *Cucurbita* genus. At the subspecies rank, three taxa are recognised: subsp. *pepo*, known only in cultivation (zucchini, pumpkins, and summer and winter squashes), subsp. *ovifera* (L.) Decker (= subsp. *texana* (Scheele) Filov), known in cultivation and in the wild (scallop and acorn squashes, ornamental gourds), and subsp. *fraterna* (L. H. Bailey) Lira, Andres & Nee (= *C. fraterna* L. H. Bailey), known only in wild populations^1–3^. Subspecies *pepo* and *ovifera* include many edible-fruited cultivar-groups, such as Pumpkin, Vegetable Marrow, Cocozelle, Zucchini, Acorn, Scallop, Straightneck and Crookneck. There is evidence of an early domestication of this species^4^, with more than one domestication event, in Mexico and United States^5^, and it has had two different diversification processes; one in America and one in Europe^6^, where Zucchini and other elongated forms, such as Vegetable Marrow and Cocozelle, were developed.

*Cucurbita pepo* is an economically important crop. Its production reached 25 million tonnes in 2014, with nearly two million cultivated hectares (http://www.fao.org/faostat/en). Cultivated varieties display a rich diversity on vine, flowering and fruit traits, and among them, cultivars of the Zucchini group rank among the highest-valued vegetables worldwide^7^. The *Cucurbita* genus and the Cucurbitaceae family contain other important crops, such as other squashes, pumpkins and gourds (*Cucurbita maxima* Duchesne and *Cucurbita moschata* (Duchesne ex Lam.) Duchesne ex Poir.), melon (*Cucumis melo* L.), cucumber (*Cucumis sativus* L.) and watermelon (*Citrullus lanatus* (Thunb.) Mansf).

Despite the agronomic importance of the species, before the genome assembly presented here, only a few *C. pepo* genetic and genomic resources were available: a first generation of genetic maps constructed with AFLP, RAPD and SSR markers^8–12^, that were later improved with SNPs^13^, and several transcriptomes^14–17^. More recently, a high density SNP based genetic map was developed using a RIL population derived from the cross between two *C. pepo* subspecies (subsp. *pepo* Zucchini × subsp. *ovifera* Scallop)^18^. This map was developed to assist us with the *de-novo* assembly process.

In the current study, we present a *de novo* assembly of the *C. pepo* genome, a high coverage transcriptome of *C. pepo*, and 40 transcriptomes of 12 species of the *Cucurbita* genus. The comparative and phylogenetic analyses show that a Whole Genome Duplication (WGD) happened just before the speciations that created this genus. All these resources and several previous transcriptome and draft genome versions are publicly available at https://bioinf.comav.upv.es/downloads/zucchini

## Material and Methods

### Plant material, genetic material isolation and NGS sequencing

Genomic DNA was isolated from nuclei of the *Cucurbita pepo* subsp. *pepo* cultivar-group Zucchini, accession BGV004370 (also referred to as MU-CU-16 and held at the COMAV-UPV Genebank, https://www.comav.upv.es). Leaves were frozen in liquid nitrogen, crushed in a mortar, and put in a solution of 0.4 mM sucrose, 10 mM Tris-HCL pH 8.0, 10 mM MgCl_2_ and 5 mM *β*-mercaptoethanol (20 ml per gram of leaves). This mixture was incubated on ice for 5 minutes. To eliminate debris and cellular fragments, samples were successively filtered through two filters (140 and a 70 *μ*m respectively), and then centrifuged at 3000 g during 20 minutes at 4 °C. The pellet was resuspended in a solution of 0.25 mM sucrose, 10 mM Tris-HCl pH 8.0, 10 mM MgCl_2_, 1% Triton X-100 and 5 mM *β*-mercaptoethanol (1 ml per gram of leaves), and centrifuged again at 12000 g for 10 minutes at 4 °C. Finally, the pellet was resuspended in 0.5 ml of 1.7 mM sucrose, 10 mM Tris-HCl ph 8.0, 2 mM MgCl_2_, 0.15 % triton X-100 and 5 mM *β*-mercaptoethanol, and then centrifuged at 18000 g during 1 hour at 4 °C. The precipitated nuclei were resuspended in CTAB buffer and the DNA was extracted using the CTAB protocol^19^. Five genomic libraries were prepared: a 500 bp pair-end library and four mate-pair libraries of 3, 7, 10 and 20 Kb insert size respectively. The first three libraries were prepared and sequenced by Macrogen (Seul, Republic of Korea) using two Illumina Hiseq2000 lanes, one for the pair-end library and another for the 3 and 7 Kb mate-pair libraries. The 10 and 20 Kb libraries were prepared by the Boyce Thompson Institute (Ithaca, New York, USA) using the Nextera protocol and were sequenced in a single Illumina Hiseq 2000 lane.

Two different sets of transcriptomes were obtained in the present study: 1) a multi-tissue transcriptome from two cultivars, representing the two main *C. pepo* subspecies to assist the genome annotation, and 2) a group of 40 transcriptomes from 12 different wild and cultivated species of the *Cucurbita* genus for the phylogenetic and comparative analyses (see Suppl. Table 1). In all cases, RNA was isolated using TRI Reagent (Sigma), treated with DNAse and purified by a chloroform and ethanol precipitation. For the *C. pepo* transcriptome, two cultivars with contrasting phenotypes were used (BGV004370 or MU-CU-16, subsp. *pepo* cultivar-group Zucchini; and BGV005203 or UPV-196, subsp. *ovifera* cultivar-group Scallop). RNA was extracted from different tissues: roots, leaves, apical shoots from plants in the male and female phase of development, flower buds collected at two early stages of flower development, mature flowers, pre-harvest fruits at different days after pollination, and post-harvest fruits subjected to different postharvest treatments (ethylene, methylcyclopropene and cold). Equivalent amounts of RNA from each tissue were mixed into two pools, one per cultivar, and two independent cDNA libraries were prepared and sequenced in an Illumina Hiseq2000 lane by Macrogen (Seul, Republic of Korea).

**Table 1.**
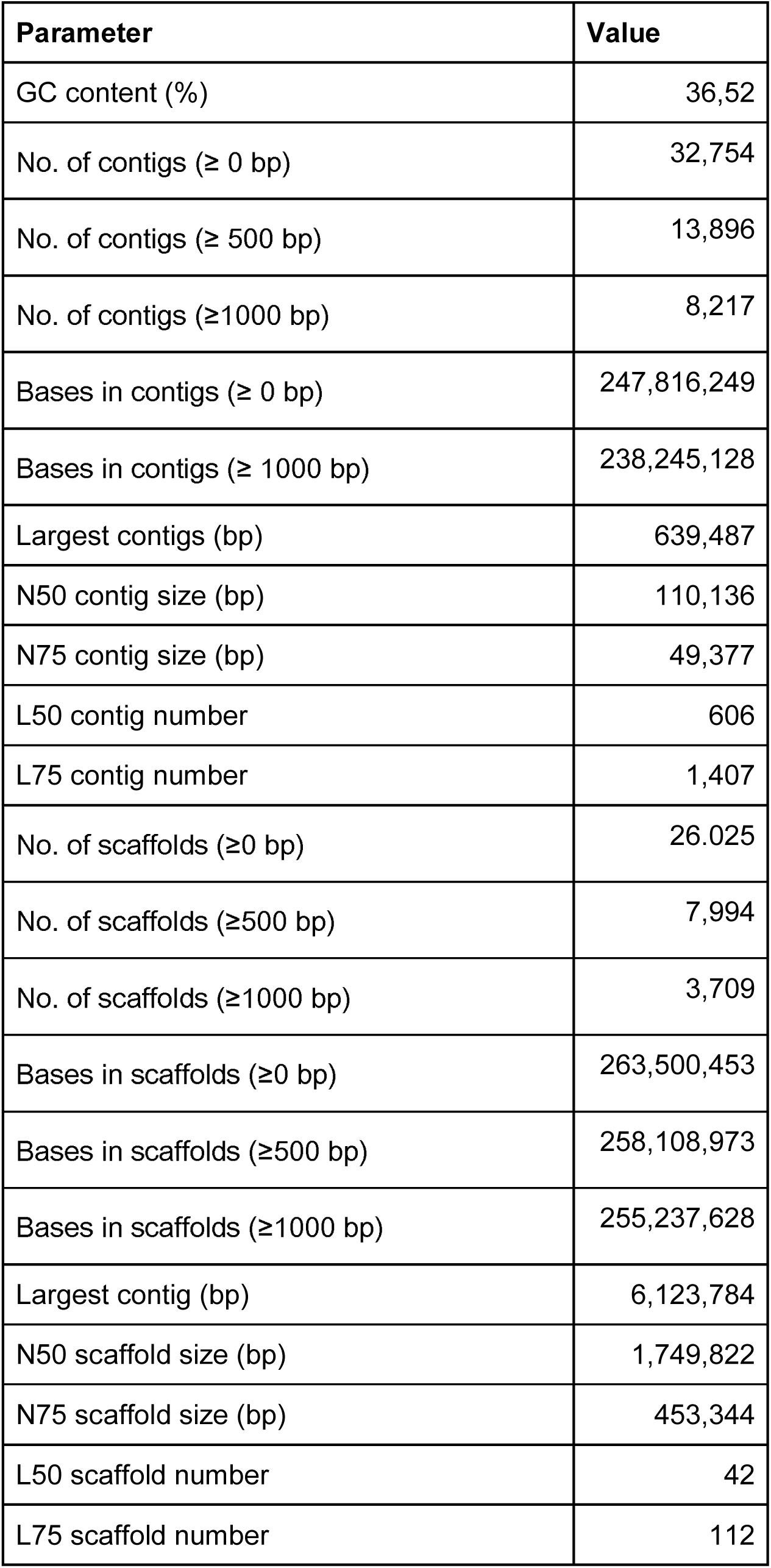
Assembly statistics of *C. pepo* genome version 4.1.

| Parameter | Value |
| --- | --- |
| GC content (%) | 36,52 |
| No. of contigs ($\geq 0$ bp) | 32,754 |
| No. of contigs ($\geq 500$ bp) | 13,896 |
| No. of contigs ($\geq 1000$ bp) | 8,217 |
| Bases in contigs ($\geq 0$ bp) | 247,816,249 |
| Bases in contigs ($\geq 1000$ bp) | 238,245,128 |
| Largest contigs (bp) | 639,487 |
| N50 contig size (bp) | 110,136 |
| N75 contig size (bp) | 49,377 |
| L50 contig number | 606 |
| L75 contig number | 1,407 |
| No. of scaffolds ($\geq 0$ bp) | 26.025 |
| No. of scaffolds ($\geq 500$ bp) | 7,994 |
| No. of scaffolds ($\geq 1000$ bp) | 3,709 |
| Bases in scaffolds ($\geq 0$ bp) | 263,500,453 |
| Bases in scaffolds ($\geq 500$ bp) | 258,108,973 |
| Bases in scaffolds ($\geq 1000$ bp) | 255,237,628 |
| Largest contig (bp) | 6,123,784 |
| N50 scaffold size (bp) | 1,749,822 |
| N75 scaffold size (bp) | 453,344 |
| L50 scaffold number | 42 |
| L75 scaffold number | 112 |

In the case of the 40 transcriptomes, the analyzed species included, besides the two *C. pepo* cultivars used in the multi-tissue transcriptome (Zucchini and Scallop), five additional genotypes of *C.pepo* (one subsp. *ovifera* (Acorn), two subsp. *pepo* (Pumpkin), and two subsp. *fraterna*). Also the four additional domesticated taxa within the species were represented: *C. moschata* (three transcriptomes), *C. maxima* (three) and its wild ancestor *C. maxima* subsp. *andreana* Naudin (South America and Africa) (one), *C. argyrosperma* Huber (Southern USA and Central America) (five), and *C. ficifolia* Bouché (Guatemala) (two), as well as six wild species occurring in Mexico and Central and South America: the mesophytic annuals *C. ecuadorensis* Cutler & Whitaker (three), *C. okeechobeensis* (Small) L.H Bailey subsp. *martinezii* (L.H.Bailey) T.C. Andres & G.P. Nabhan ex T.W. Walte (three), and *C. lundelliana* L.H Bailey (four) and the xerophytic perennials *C. foetidissima* Kunth (four), *C. cordata* S.Watson (two) and *C. pedatifolia* L.H.Bailey (three). RNA was extracted exclusively from young leaves and the cDNA libraries were prepared and sequenced in a Hiseq2000 lane in the Boyce Thompson Institute (Ithaca, New York, USA).

### *De-novo* genome assembly

The pair-end and mate-pair reads were cleaned using the *ngs_crumbs* software (code available at https://github.com/JoseBlanca/) to eliminate adapters, low quality bases (Phred quality < 25 in a 5 bp window), reads shorter than 50 bp, and duplicated sequences. The Nextera mate-pair reads (10 Kb and 20 Kb libraries) were classified by NextClip v0.8^20^ according to the presence of the junction adaptor. Only the mate-pairs in which NextClip was able to detect and trim the adaptor were used for the assembly. For the pre-Nextera mate-pair libraries, the detection and filtering of possible chimeric pairs was done by mapping the reads against a first assembly of the genome and only the pairs with the expected orientation and at the expected distance were kept. The implementation of this process can be found in the *classify_ chimeras* and *trim_ mp_chimeras* binaries of the *ngs_chrumbs* software. The mitochondrial and chloroplastic reads were detected by blasting^21^ them against the *C. melo* organelle genomes (JF412791.1 and NC014050.1). Mitochondrial and chloroplastic reads were also included in the assembly, but only enough randomly selected reads to get a 150X coverage. Assemblies with k-mer lengths from 31 to 61 with a step-size of 4 were carried out. The final assembly was done by SOAPdenovo2 v2.04^22^ using k-mer size of 41. Resulting scaffolds were broken with BreakScaffolds (https://github.com/aubombarely/GenoToolBox) and reassembled with SSPACE^23^. The new scaffolds were improved using SOAPdenovo2’s GapCloser^22^. Gene completeness of the assembly was assessed using BUSCO v.2^24^. Mitochondrial and chloroplastic scaffolds were identified using BLAST^21^ against the chloroplast and mitochondrial genomes of *C. melo*. Genome size was estimated from the k-mer depth distribution as ∑(*d* · *k_d_*) / D where *d* is the k-mer depth, *k_d_* is the number of k-mers for the given depth and *D* is the maximum k-mer depth of the distribution. The leftmost part of the distribution was discarded as it includes mostly k-mers due to sequencing errors. The k-mer distribution was calculated by Jellyfish^25^ using a k-mer size of 31.

In order to detect assembly artifacts and to group scaffolds into pseudomolecules, a genetic map was built. A group of 120 individuals of a F_8_ Recombinant Inbreed Line (RIL)^18^, developed through single seed descent from a previous Zucchini (BGV004370) x Scallop (BGV005203) F_2_^13^, were genotyped by Genotyping-by-sequencing (GBS)^26^. SNP calling was performed using Freebayes^27^ and a genetic map was constructed using the R packages R/qtl^28^ and ASMap^29^ (see details in Montero-Pau et al.^18^). Scaffolds that were present in more than one linkage group in the genetic map were visually explored with Hawkeye^30^ and manually splitted. Scaffolds were ordered and oriented according to the genetic map into pseudomolecules.

### *De-novo* transcriptome assembly

Raw reads were processed using *ngs_crumbs* software to eliminate adapter sequences, low quality bases (Phred quality < 25 in a 5 bp window) and sequences shorter than 40 bp. The transcriptome was assembled with the Trinity assembler v2.0.6^31^ with default parameters. In the case of the *C. pepo* transcriptome, reads of both cultivars were merged in order to get a more comprehensive representation of the transcriptome. Additionally, reads from a previous 454-based transcriptome^17^ were also included. The resulting contigs were reassembled with CAP3^32^ to eliminate redundancies. Low complexity transcripts were filtered out using *ngs_crumbs*. Trinity subcomponents were clustered using BLAST into unigene clusters doing a transitive clustering. Any two transcripts that shared an overlap longer than 100 bp and a similarity higher than 97% were considered to belong to the same unigene cluster. Finally, transcripts expressed less than 1 % of the most expressed transcript in each Trinity subcomponent were filtered out using RSEM (http://deweylab.biostat.wisc.edu/rsem/).

### Genome annotation

Genome structural annotation was performed using Maker-P^33^ (version 2.31.6) with the default parameters. The *C. pepo* transcriptome was used to train Augustus^34^ (version 3.0.2) with the default parameters. SNAP^35^ (version 2006-07-28) was also trained with the same dataset following the instructions from the Maker-P manual. Repetitive sequences were extracted from the genome reference using RepeatModeler^36^ (version 1.0.8). The *C. pepo* transcriptome, repetitive sequences, and training ab-initio gene predictor files were used for the annotation with Maker-P. Functional annotation was performed by sequence homology search using BlastP (minimum E-value of 10^−10^) with GenBank, TAIR10 and SwissProt protein datasets (downloaded 2014-12-21). Additionally, InterProScan^37^ was used to annotate protein domains, extending the annotation to Gene Ontology terms associated with these protein domains. Blast2GO^38^ was used to do an annotation based on a Blast search against NCBI’s nr database. Functional descriptions were processed using AHRD (https://github.com/groupschoof/AHRD) giving a weight of 100, 50 and 30 to SwissProt, TAIR and GenBank annotation respectively.

A structural and homology-based approach, as described in Campbell et al.^33^, was used to annotate the repetitive DNA. Briefly, miniature inverted repeat transposable elements (MITE) and long terminal repeat (LTR) retrotransposons were collected using MITE-Hunter^39^, LTR-harvest, and LTR-digest40,41. A MITE and LTR library was built after excluding false positives, and selecting representative sequences^42^. This library was used to mask genome sequences with RepeatMasker^36^, and the resulting sequences were then processed by RepeatModeler in order to look for other repetitive sequences.

Reference sequences of Copia and Gypsy LTR superfamilies of the retrotranscriptase gene were obtained from GyDB^43^. Sequences were manually aligned, and best fitting nucleotide substitution model (based on Bayesian information criterion) and maximum-likelihood tree for each superfamily were obtained using IQ-TREE^44,45^. Branch support was computed using the bootstrap ultrafast method.

### Transcriptome annotation

Transcripts were blasted against Swiss-Prot, UniRef90, and the *Arabidopsis* proteins. Orthologues with cucumber and *Arabidopsis* were detected using a bi-directional BLAST search. The unigenes were associated to GO terms using Blast2GO software^38^. ORFs were predicted in the unigenes with the aid of the ESTScan software^46^.

### Comparative genomics

Four complete genomes of three related species belonging to the Cucurbitaceae family were included in the study for comparative genomic analyses: *Citrullus lanatus* (genome v. 1)^47^, *Cucumis sativus* var. *sativus* (Chinese long) (v. 2)^48^, *C. sativus* var. *hardiwickii* (Royle) Gabaer (PI 183967) (v. 1) and *Cucumis melo* (v. 3.5)^49^. The first three are accessible at www.icugi.org and the later at http://melonomics.net. In order to be able to compare among genomes, the repetitive DNA characterization previously described was performed in these four genomes.

Detection of gene duplications were carried out in the gene families created by using OrthoMCL^50,51^ and OrthoMCL DB version 5 on the predicted proteomes of the five cucurbit genomes. In those cases in which more than one transcriptional variant was found for the same gene, only the longest variant was used. Differences in the functional role of the duplicated genes were assessed through GO enrichment tests using R package topGO^52^, and REVIGO^53^ was used to visualize the results. Rate of transversions on 4-fold degenerate synonymous sites (4DTv) was calculated between pairs of orthologs and paralogs using an in-house Python script. Values were corrected for multiple substitutions^54^.

The phylogeny (40 transcriptomes and 5 genomes) was reconstructed using a concatenated method and through a joint estimation of both species and gene trees carried out by Phyldog^55^. For the first approach, single copy genes detected using OrthoMCL that were present in all cucurbit genomes were selected, and then the corresponding *C. pepo* transcript was blasted against the 40 *Cucurbita* spp. transcriptomes. Only the blast hits with an E-value higher than 10^−60^ and a match longer than 200 bp were retained. For each gene family, sequence alignments were built using an iterative refinement method implemented in MAFFT^56^. Alignments with less than 30 species were excluded. All resulting gene families were concatenated and the maximum-likelihood tree was inferred using IQ-TREE^44^ using a nucleotide substitution model for each gene^45^. For each partition, the best model was selected based on the Bayesian information criterion (BIC). Branch support was obtained by bootstrap using an ultrafast method^57^.

For the Phyldog approach sequences were clustered in ortholog groups by blasting all *C. pepo* genes against all 40 *Cucurbita* spp. transcriptomes and the four cucurbit genomes. Blast hits with an identity lower than 70% and shorter than 200 residues were ignored. For each group, three multiple sequence alignments were obtained using Kalign^58^, MUSCLE^59^ and MAFFT^56^. Alignment results were combined and evaluated with T-Coffee^60,61^ and only alignments with an alignment score higher than 900 were kept. For each alignment a starting tree for Phyldog was inferred using PhyML^62^ assuming the best nucleotide substitution model obtained by jModeltest^63^. Phyldog^55^ was then used to simultaneously infer species and gene trees and to detect gene duplication events.

## Results

### *De novo* genome assembly

The complete genome of *Cucurbita pepo* has been sequenced using a whole genome shotgun sequencing approach. The Zucchini type (*C. pepo* subsp. *pepo*) accession MU-CU-16 was selfed 4 times before sequencing. This accession is characterized by early flowering, bushy growth habit, high production, and uniform cylindrical dark green fruits. This accession was also used as parental in two previous genetic maps^13,18^. One paired-end, with an insert size of 500 bp, and four mate-pair, with sizes of 3, 7, 10 and 20 Kb, libraries were created and sequenced in 5 Illumina Hiseq2000 lanes, resulting in a genome coverage of 254 X for the pair-end library and 54, 46, 65 and 62 X for the 3, 7, 10 and 20 Kb libraries, respectively (Suppl. Table 2). All reads were quality trimmed and filtered. Additionally, about 40% of the 3 and 7 Kb mate-pair reads were found to be chimeric and filtered out by comparing them against a preliminar assembly (see Suppl. Table 2, Suppl. Fig. 1). This chimeric filtering doubled the contig N50 and tripled the scaffold N50 of the final assembly. Finally, 503 M filtered pair-end reads and 185 M mate-pair reads were used in the assembly. The genome was assembled by SOAPdenovo2^22^. A k-mer size of 41 was chosen for the final assembly because it rendered the highest N50 values (Suppl. Fig 2). The SOAPdenovo2 scaffolds were broken and the scaffolding was redone with SSPACE^23^ and GapCloser^22^. The final assembly covered 263 Mb in 26,005 scaffolds and 32,754 contigs with a contig N50 of 110 Kb (L50 = 606 contigs) and a scaffold N50 of 1.8 Mb (L50 = 42 scaffolds) (Table 1). Completeness of the *de novo* assembly was assessed with BUSCO using a plant-specific database of 1440 genes. 92.1% of them were found complete (73.1% as single genes and 19.0% as duplicated) and 2.1% were found fragmented. The Illumina RNAseq reads obtained from the MU-CU-16 accession were mapped with HISAT2 with this genome as reference with a 91.9% success rate. The pair-end reads used to build the assembly were mapped against the assembly with a success rate of 99.4%. From the k-mer distribution genome size was estimated to be 283 Mb, thus 93.0% of the genome would be covered by the assembly. Chloroplastic and mitochondrial scaffolds were detected using Blast: 250 scaffolds were identified as mitochondrial and 13 as chloroplastic (Suppl. Table 3).

**Figure 1.**
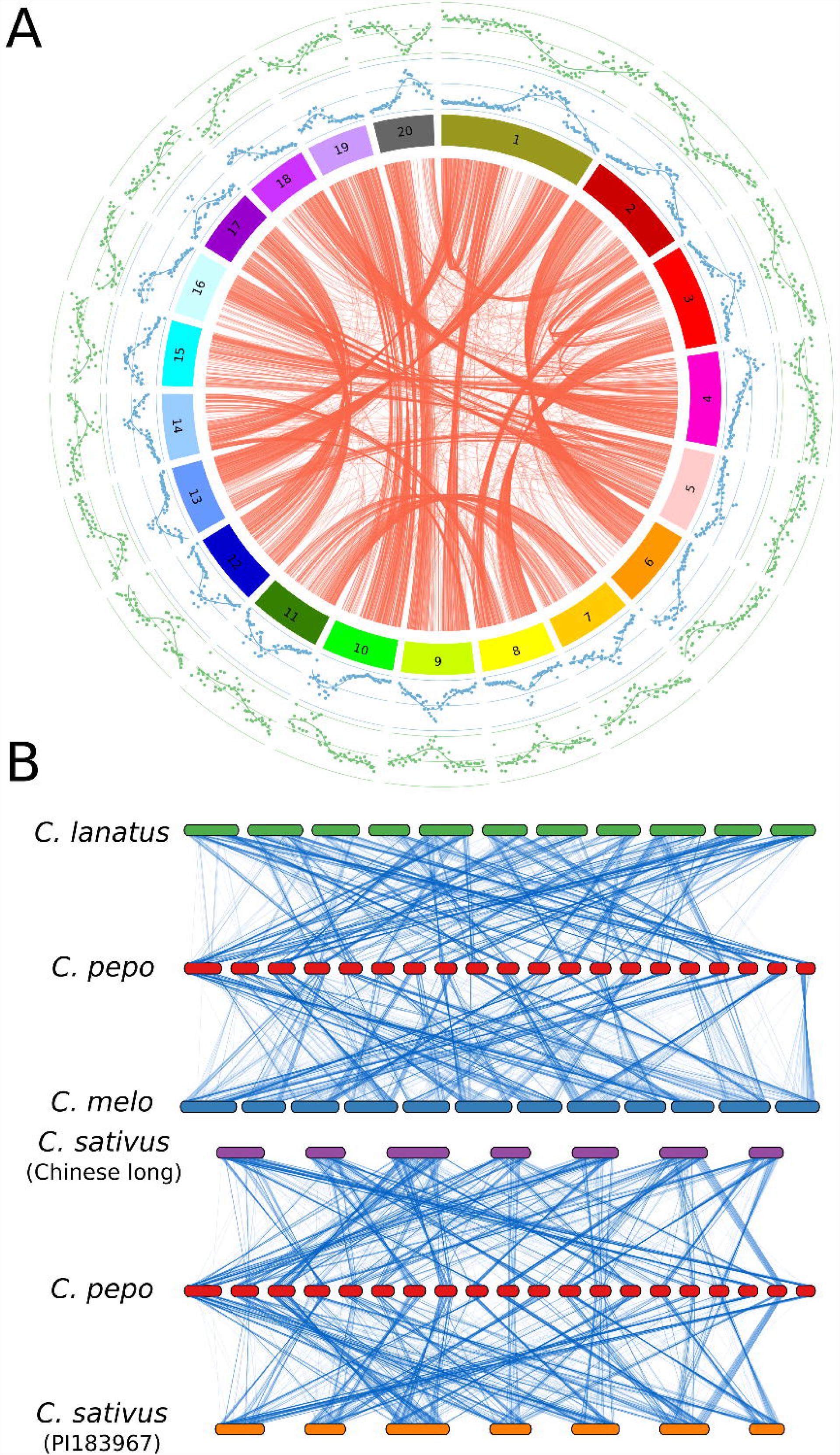
Genome organization. A) Circos plot showing paralog gene pairs in *Cucurbita pepo* (red lines). Outer plots represent the proportion of repetitive (blue) or gene coding (green) DNA by 200 Kb windows. B) Genomic synteny between *Cucurbita pepo* and *Cucumis melo, Cucumis sativus* and *Citrullus lanatus.* Lines join single copy orthologs.

**Figure 2.**
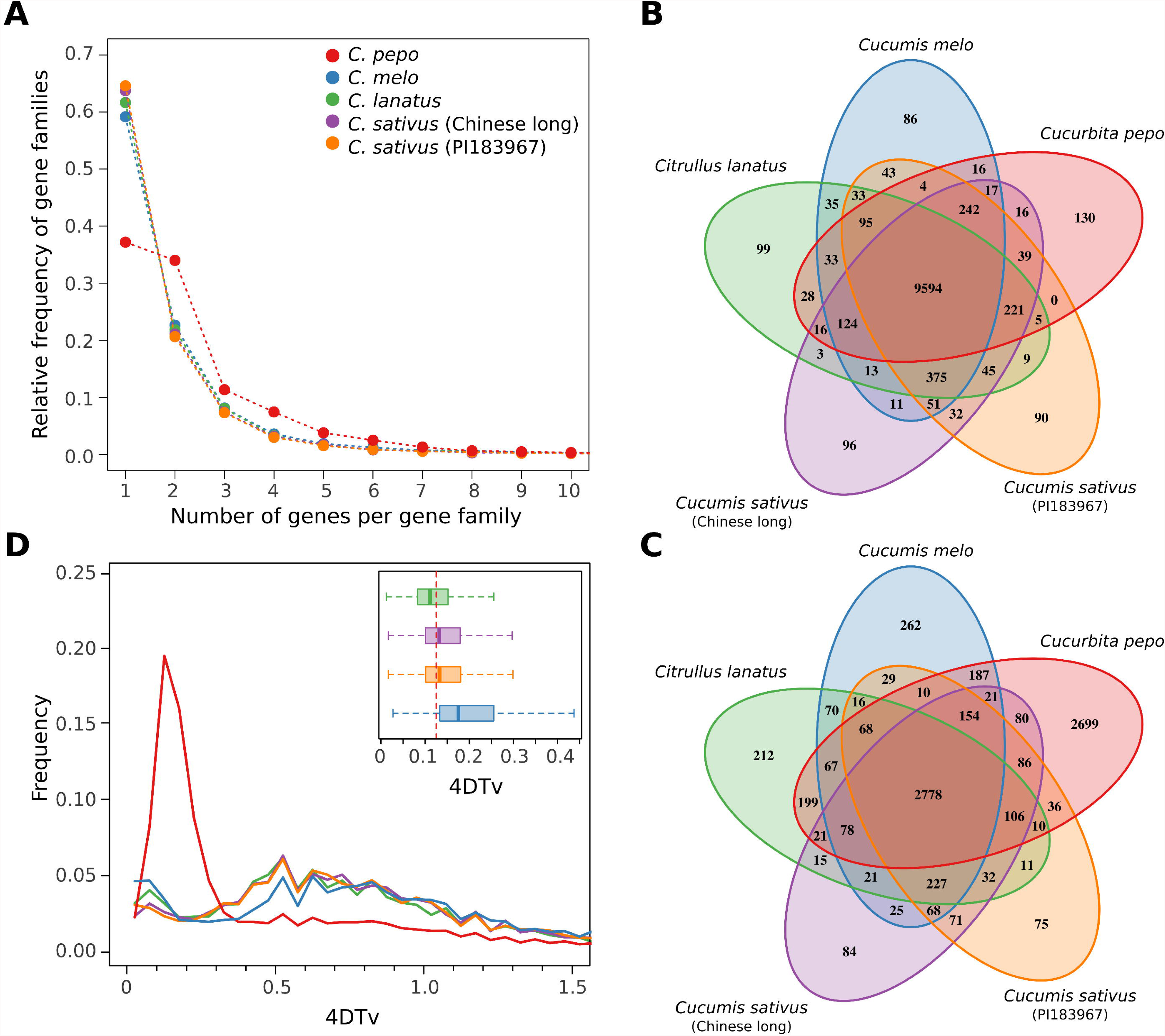
Genome duplication. A) Distribution of the number of gene families based on the number of gene copies for *Cucurbita pepo, Cucumis melo, Cucumis sativus* and *Citrullus lanatus*; B) Venn diagram showing the number of gene families, and C) the number of duplicated gene families shared among the cucurbit genomes; D) distribution of the rate of transversions on 4-fold degenerate synonymous sites (4DTv) among paralogs for the five studied genomes. Inset shows the boxplots for the 4DTv distribution between ortholog copies of *C. pepo* and the rest of cucurbit species. Red dashed line shows the duplication event in *C. pepo.*

**Table 2.** Pseudochromosome summary. Number of scaffolds anchored to each pseudochromosome, total length and length without the 1000 N spacers.

| Molecule | #Scaffolds | Length (bp) | Length without N spacers (bp) |
| --- | --- | --- | --- |
| Cp4.1LG01 | 19 | 21,320,769 | 21,302,769 |
| Cp4.1LG02 | 16 | 14,376,414 | 14,361,414 |
| Cp4.1LG03 | 12 | 13,772,414 | 13,761,414 |
| Cp4.1LG04 | 5 | 12,709,140 | 12,705,140 |
| Cp4.1LG05 | 8 | 10,865,678 | 10,858,678 |
| Cp4.10LG06 | 11 | 10,677,745 | 10,667,745 |
| Cp4.1LG07 | 14 | 10,147,556 | 10,134,556 |
| Cp4.1LG08 | 4 | 10,059,303 | 10,056,303 |
| Cp4.1LG09 | 10 | 9,920,322 | 9,911,322 |
| Cp4.1LG10 | 8 | 9,835,092 | 9,828,092 |
| Cp4.1LG11 | 11 | 9,833,969 | 9,823,969 |
| Cp4.1LG12 | 5 | 9,824,194 | 9,820,194 |
| Cp4.1LG13 | 8 | 9,354,089 | 9,347,089 |
| Cp4.1LG14 | 5 | 8,955,933 | 8,951,933 |
| Cp4.1LG15 | 4 | 8,816,444 | 8,813,444 |
| Cp4.1LG16 | 10 | 8,691,934 | 8,682,934 |
| Cp4.1LG17 | 9 | 8,680,504 | 8,672,504 |
| Cp4.1LG18 | 7 | 8,333,454 | 8,327,454 |
| Cp4.1LG19 | 8 | 8,246,682 | 8,239,682 |
| Cp4.1LG20 | 7 | 8,120,804 | 8,114,804 |

**Table 3.** Genome annotation summary.

|  |  |
| --- | --- |
| Genes | 34,240 |
| Protein coding genes | 27,870 |
| mRNAs | 27,870 |
| Protein-coding mRNAs | 27,870 |
| Exons | 184,243 |
| CDSs | 166,271 |
| Introns | 150,003 |
| 5' UTRs | 21,701 |
| 3' UTRs | 22,296 |
| tRNAs | 6,370 |

The genetic map developed with the RIL population of Zucchini x Scallop (accessions MU-CU-16 x UPV-196)^18^ was used to detect chimeric scaffolds and to anchor and order the scaffolds into pseudomolecules. 7,718 SNPs (average of 386 markers/linkage group) were located in the map. Based on the relationship of physical and genetic distances and on the presence of the same scaffold in more than one linkage group, 22 out of the 26,005 scaffolds were identified as chimeric. Those scaffolds were visually inspected and splitted. In a first attempt, a total of 181 scaffolds could be anchored to 21 pseudomolecules, which represents the 81,4% of the assembled genome. Finally, after the integration of this genetic map with the genetic maps developed by Esteras et al.^13^, that was based on data from the F_2_ of the same cross, and the genetic map of Holdsworth et al.^64^, all scaffolds were grouped into 20 pseudomolecules (Table 2 and Suppl. Table 4). Between 4 and 19 scaffolds were anchored to each pseudochromosome with a length between 8.1 Mb and 21.3 Mb (Table 2). Out of the remaining 25,344 scaffolds, 3,295 were longer than 1Kb and 365 were longer than 20Kb. The average correlation between physical distance and genetic distance was 0.98 (0.94-1.00) (Suppl. Fig. 3). This assembly constitutes genome version 4.1. Some other previous versions were made available to the *Cucurbita* community, but none were published.

**Figure 3.**
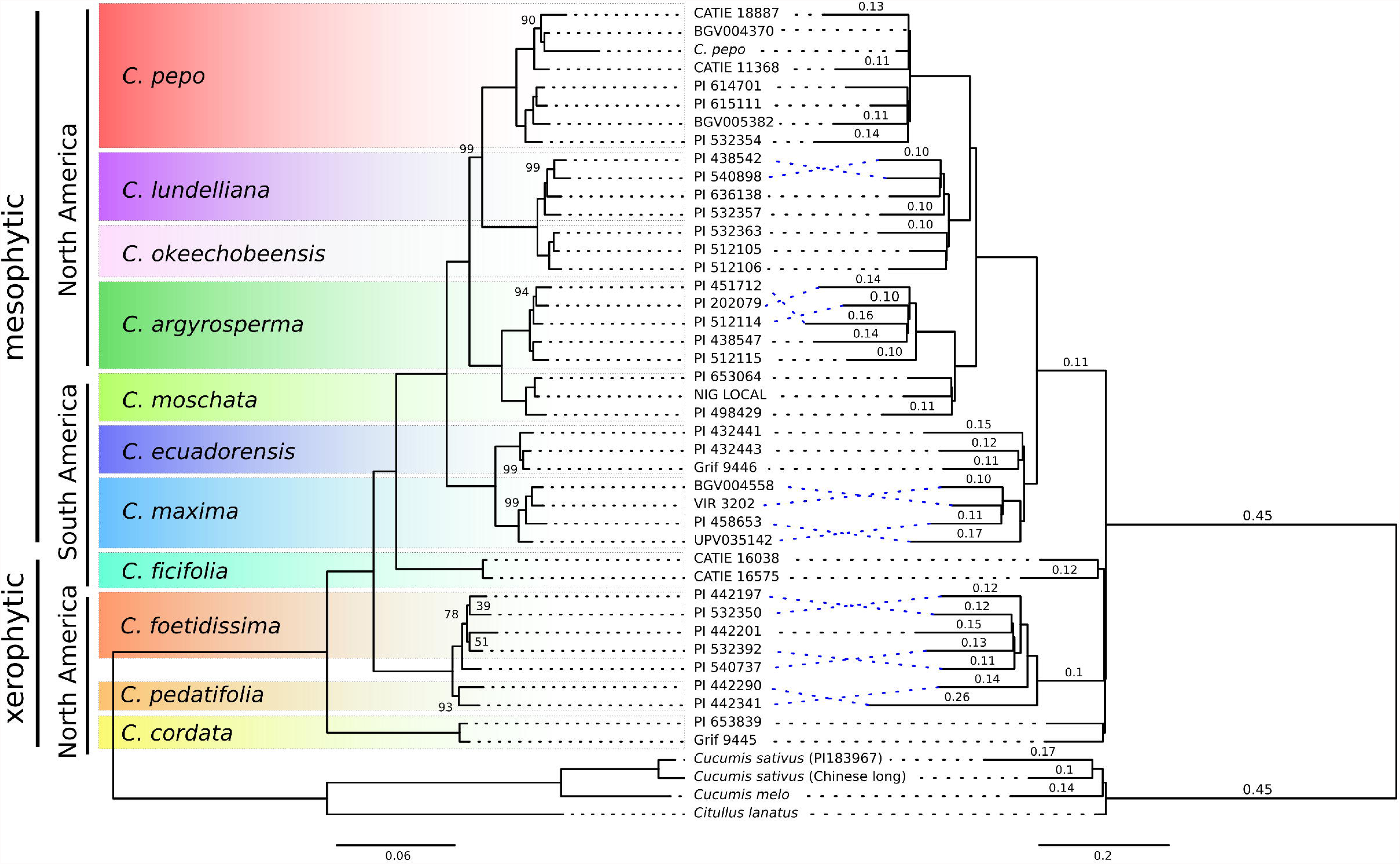
Phylogeny of *Cucurbita* genus based on a concatenated method (left tree) or a joint estimation of gene and species trees (right tree). Left tree: branch lengths represent genetic distance and only bootstrap values lower than 100 are showed. Right tree: branch length represent proportion of duplicated genes per branch, values shown those proportions of duplicated genes higher than 0.1

### Transcriptome and genome annotation

Two cDNA libraries were created for the parent accessions of the RIL population using pooled RNA from different vegetative and reproductive tissues. More than 228 millions of reads were added to the previously available 454-based transcriptome^17^. They were used to create a new transcriptome assembly (version 3.0, available at https://bioinf.comav.upv.es/downloads/zucchini) and to annotate the genome. The transcriptome assembly identified 108,062 transcripts, 65,990 of which included an ORF. GO terms could be assigned to 71.5% of the coding transcripts.

The genome annotation resulted in 34,240 predicted gene models, out of which 27,870 were protein-coding genes (Table 3). These results are similar to those found in melon and cucumber^48,49^. The average gene size was 3,450 bp with an average number of exons of 5.4 (Suppl. Fig. 4). The gene models cover 118 Mb, and their coding regions 35 Mb, which represents 45.3% and 13.7% of assembled genome respectively, and indicates a high degree of genome compaction (Fig. 1A). GO terms could be assigned to 19,784 protein-coding genes out of 27,870 (71,0%) (Suppl. Fig. 5 and Suppl. Fig. 6). Functional descriptions were added to 76.6% of transcripts using AHRD, and 79.2% were tagged with an IntrePro protein domain.

### Repetitive elements

We identified that 93 Mb (37.8% of the assembly) consisted of repetitive elements (REs) (Suppl. Fig. 7 and Suppl. Table 5). Long terminal repeats (LTR) represented 50.7% of the identified REs. *Gypsy* and *Copia* were the most abundant LTR superfamilies (24.2% and 19.8% of identified REs, and 3.3% and 2.7% of the total genome). The *Gypsy* LTR abundance is similar to that found in *C. melo, C. lanatus* and *C. sativus,* which ranged from 19.5 to 34.4%, whereas the *Copia* family was less represented than in other Cucurbitaceae genomes (30.9 - 34.4%). Other two LTR superfamilies were more copious in the *C. pepo* genome than in the other cucurbits: *Cassandra* (3% of identified RE vs. 0.1 - 0.8%), and *Caulimovirus* (2.1% vs. 0.26- 0.9%). Satellites and simple repeats constituted 25.2% of all identified REs, which is a larger fraction than in related Cucurbitaceae species (4.4% - 12.4%). *Copia* and *Gypsy* REs were assigned to their different families by building two phylogenetic trees (one for *Copia* and one for *Gypsy*) (Suppl. Fig. 7). All *Copia* and *Gypsy* families previously identified in *C. melo, C. lanatus* and *C. sativus* were also present in *C. pepo,* except for *Copia/Bianca* and *Gypsy/Ogre* families. In these trees the Gypsy/*Galadriel* and *Copia/Tork4* families were overrepresented in *C. pepo*, so they seem to have suffered a diversification process in this species. Finally, approximately, 24% of REs were not assigned to any class of repetitive or transposable elements (TE).

### Comparative genomics

Genes, represented by its longest protein, of the four cucurbit crops: *Cucurbita pepo*, *Cucumis melo, Citrullus lanatus*, and two *Cucumis sativus* cultivars (var. *sativus,* Chinese long; and var. *hardiwickii,* PI 183967) were grouped into gene families using OrthoMCL. The percentage of genes that could be assigned to a gene family in these species ranged from 91.2 to 72.8% (Suppl. Table 6). In *C. pepo* the number of gene families with two or more paralogs was higher than in the other crops (Fig. 2 A). Most *C. pepo* gene families were also present in the other cucurbits (Fig. 2 B), however many of them had more than one gene in *C. pepo* (Fig. 2 C). Most of the Zucchini paralogs were organized in large syntenic regions that cover most of the genome (Fig. 1). Synteny with the other cucurbit species showed that despite some conserved synteny, an extensive chromosomal rearrangements has occurred (Fig. 1). The high number of paralogous genes detected and their synteny suggests that *C. pepo* could have suffered a WGD (Suppl. Tables 7, 8 and 9).

The rate of transversions on 4-fold degenerate synonymous sites (4DTv) is a neutral genetic distance that can be used to estimate relative timing of evolutionary events. The distribution of 4DTvs among paralog pairs for all species, but *C. pepo*, showed a wide peak that ranged from 0.4 to 1.1 with a maximum about 0.6 (Fig. 2 D), whereas for *C. pepo* a more recent and narrower peak centered around 0.12 was found. Speciation can also be relatively dated by computing the 4DTvs between orthologous genes of any pair of species. This showed that the speciation event that gave rise to the *Cucurbita* genus, represented by the pairwise 4DTv distributions of *C. pepo* against *C. lanatus, C. sativus,* and *C. melo,* occurred almost simultaneously with the duplication event found in *C. pepo* (Fig. 2 D).

A total of 40 transcriptomes were assembled by Trinity from Illumina reads for 12 species, this resulted in 18,446 to 67,366 genes and 18,902 to 92,522 transcripts (Suppl Table 1). The species and gene family trees were reconstructed using Phyldog, including both the genomes and these 40 transcriptomes. Phyldog marked the duplication events in the gene family trees and calculated the number of duplications per branch in the species tree. According to Phyldog most gene families suffered a duplication event (90%) in the branch that originated the *Cucurbita* genus (Fig. 3). Additionally, a maximum likelihood phylogeny was reconstructed using a concatenated alignment using IQ-TREE. The topologies recovered by both methods are highly congruent, the species trees based on genomic data, showed that xerophytic perennial species (*C. cordata*, *C. pedatifolia*, and *C. foetidissima*) were in a basal position, while mesophytic annuals or short-lived perennials species of the genus were derived from them and formed a monophyletic taxon. The only remarkable difference between both methods was the position of *C. ficifolia,* IQ-TREE grouped it with the mesophytic species, whereas the Phyldog tree grouped it with the xerophytic species. There are some other minor differences related with the position of some accessions within a particular species between both trees. Some of these differences are related to suspected hybrid accessions like PI540737 (between *C. pedatifolia* and *C. foetidissima*) or PI532392 (between *C. scabridifolia* and *C.foetidissima*). In general, all nodes are supported by bootstrap values close to 100 except those related with hybrids.

A GO enrichment analysis was carried out on three sets of genes: 1) single copy *C. pepo* genes, 2) all duplicated *C. pepo* genes, and 3) duplicated *C. pepo* genes found to be single copy in the rest of cucurbits (i.e., melon, watermelon and cucumber). Single copy genes were enriched in nucleic acid metabolic processes, DNA repair, DNA replication, DNA recombination, rRNA and tRNA processing, lipid metabolism and embryo development (Suppl. Table 10 and Suppl. Fig. 8). In the gene set found to be duplicated in all species, the most significantly enriched GO terms were: transcription and translation regulation, protein metabolism, transmembrane transport, ribosome biogenesis and signal transduction. The terms enriched in the *C. pepo* exclusive duplication were: NAD biosynthesis, regulation of signal transduction, mitochondrial respiratory chain, regulation of cell cycle and cell structure, intracellular protein transport, pollen and vegetative development, photosynthesis light harvesting, and photoperiodism flowering. Interestingly, other genes related to flower development were also found among the exclusively duplicated genes in *C. pepo* such as EARLY FLOWERING 4, Zinc finger CONSTANS-LIKE 3, flowering locus T, RTF1, CDF73, KNUCKLES, CTR9, flowering locus K, GID1b, FLC EXPRESSOR, FRIGIDA and FPA. Additionally, seven out of 34 genes annotated as “similar to CONSTANS-LIKE protein” were exclusively duplicated in *C. pepo,* as well as five out of 9 genes annotated as “similar to FRIGIDA”.

## Discussion

In this study we present the first description of the *C. pepo* genome. This new assembly is organized in 20 pseudomolecules, has a scaffold N50 of 1.8 Mb, and is integrated with a high density genetic map. According to the coverage (92.1%) of the BUSCO conserved gene core set and the percentage of the RNAseq reads (91.1%) and genomic reads (99.4%) mapped against it, the current assembly covers most of the zucchini genome. The genome size inferred by k-mer analysis was 283 Mb, so this assembly would constitute 93.0% of the genome. Thus, this assembly is an almost complete representation of the *C. pepo* genome.

Our results show that *C. pepo* genome has suffered a WGD that took place in the origin of the *Cucurbita* genus. Three independent evidences support this WGD: the topology of the gene family phylogenies, the karyotype organization, and the distribution of 4DTv distances. Phyldog reconstructed the phylogeny of every gene family and by comparing it with the species tree topology inferred where the duplication event likely occurred in each family. According to this analysis most duplications happened in the branch that separated the *Cucurbita* genus from the rest of species in the Cucurbitaceae family. The genome structure shown by the physical location of the pairs of Zucchini paralog genes was characterized by large syntenic regions within this species. These syntenic regions cover most of the genome and are likely to have been generated by a pseudodiplodization process of an ancestral tetraploid followed by different chromosomal rearrangements. Interestingly, all species of the *Cucurbita* genus (tribe *Cucurbiteae*) present n=20 chromosomes^65,66^, whereas species of *Benincaseae* tribe, which include *Cucumis* and *Citrullus* genera, have a different chromosomal organization with n=12 (melon), n=11 (watermelon) or n=7 (cucumber)^66^. A possible poliploidy in the origin of *Cucurbita* was already proposed based on the chromosome number and the number of isoenzyme copies^67,68^. Despite the WGD, the size of the Zucchini genome is similar to that of the other sequenced cucurbits, and also, the number of genes is not much higher. This suggests that most genes were deleted after the WGD event. It might well be the case that there is a selective pressure to keep the genome size of these species within a certain range and that the maintained genes were specifically selected.

The 4DTv distribution found in *C. melo, C. sativus* and *C. lanatus* showed no evidence of a recent WGD^48,49^. These three species present a peak on the 4DTv distribution around (0.6) that corresponds to the ancestral paleohexaploidy (γ) event that happened in the divergence of monocotyledons and dicotyledons (~ 300 Mya)^69^. However, the 4DTv distances between paralog genes within Zucchini showed lower distances characterized by a mode of 0.12. Thus most paralogs seem to have been created by a recent duplication. Additionally, the 4DTv peaks found in the distribution calculated for the orthologous genes between *C. pepo* and melon, cucumber and watermelon can be used to date the Zucchini duplication. These peaks are all very close to the Zucchini duplication peak. Thus, both the 4DTv and gene family phylogenies are consistent with a duplication that happened in the ancestral species that gave rise to the *Cucurbita* genus short after its split from the ancestor of *C. melo*, *C. sativus* and *C. lanatus* about 30 ± 4 My^70^. The evolutionary rate derived from this time estimation is consistent with that found in other plants with a recent WGD like *Nelumbo nucifera* (4DTv = 0.17, 18 Mya)^71^, *Glycine max* (0.057, 13 Mya)^72^, *Zizania latifolia* (0.07, 13 Mya)^73^, and *Setaria italica* (0.38, 70 Mya)^74^. *Populus trichocarpa* would be an exception with a much lower evolution rate (0.1, 60-65 Mya)^75^, but this discrepancy could be due to the longer generation time of this plant^76^. This WGD might have provided the *Cucurbita* species a way to generate new gene functions used to adapt to new habitats. In fact, this genus includes both xerophytic species, perennials adapted to dry climates and species adapted to moister or mesophytic environments, either annuals or short-lived perennials, and expands from tropic to template regions of America. For instance, the duplication of the photosynthesis and flower development regulation genes found in the GO enrichment analysis, could have provided mechanisms to adapt flowering to variation in temperature and the duration of days found from Southern USA to Southern South America. Genes of these pathways have been implicated in the adaptation of several crops to different photoperiod and geographical adaptation^77^. FLOWERING LOCUS T has been described as a possible long-distance florigenic signal in the cucurbits^78^. The genus also includes an amazing variation in morphological traits related to vine, fruit and seeds.

In agreement with previous *Cucurbita* phylogenetic studies^79–84^, the xerophytic perennial species (*C. cordata*, *C. pedatifolia*, and *C. foetidissima*) were basal to the *Cucurbita* genus. The current analysis supports the relationship among mesophytic species found by Kates et al.^81^, and additionally, it clarifies the clustering of the sister species *C. foetidissima* and *C. pedatifolia*, and *C. lundelliana* and *C. okeechobeensis* that were not previously resolved^81^. The position of *C. ficifolia* remains controversial. The concatenated method clusters it as a basal species to the annual mesophytic taxa, showing a paraphyletic relationship with respect to the perennial taxa, in agreement with Wilson et al.^82^ and Kates et al.^81^. However, based on Phyldog, *C. ficifolia* appears as a sister species of *C. pedatifolia* and *C. foetidissima*, in agreement with Zheng et al.^83^. *Cucurbita ficifolia* is a mesophytic species, but shares some morphological features with the xerophytic species. More data is needed to establish the relationship of *C. ficifolia* to the mesophytic/xerophytic species of the genus. Also, this incongruence between trees may be also due to hybridization, as some partially fertile hybrids have been obtained between *C. ficifolia* and *C. lundelliana*, *C. foetidissima* and *C. pedatifolia*^85^, or it might be the result of very close speciation events.

This genome assembly constitutes a key resource for the study and breeding of the economically important *C. pepo*. Previous unpublished drafts, made available by us, of this genome have already been used in several publications related to the detection of resistance genes, the study of fruit development or the generation of molecular marker sets^15,64,86^. Additionally, we have assembled 40 transcriptomes for 11 species of the *Cucurbita* genus, which can be a valuable source of molecular markers, as well as the foundation of comparative genomic studies.

## Acknowledgements

The authors want to thank the USDA, CATIE, VIR and COMAV-UPV genebanks for providing some of the accessions used in this paper. Authors also thank Cristina Roig, Gorka Perpiña and Eva Maria Martínez for their technical assistance.

## Author Contributions Statement

JB, JM-P, BP and JC designed and conceived research. JB, PZ, CM and JC contributed to the assembly of the genome and transcriptomes. JM-P, AB and LM realized the annotation of the genome. JM-P integrated genome assembly with genetic maps and analyzed genome duplication studies. BP, CE, MF selected/provided plant materials and maintained all living materials. BP and MF did species classification. JC and CE prepared the DNA samples. JC, CE, MJ and PG participated in the preparation of RNA samples for the libraries. JM-P, JB, BP and JC wrote the manuscript. All authors read and approved the final manuscript.

## Additional Information

This work was partially funded by the INIA project RTA2011-00044-C02-2 with contributions of E-RTA2013-00020-C04-03 of the Spanish Instituto Nacional de Investigación y Tecnología Agraria y Alimentaria (INIA) cofunded with FEDER funds (EU), and AGL2014-54598-C2-1-R of the Spanish Ministry of Economy and Competitivity.

Genome assembly and raw sequences are deposited in NCBI under BioProject PRJNA386743. Genome v. 4.1, genome annotation and transcriptome v. 3.0 are also available at http://bioinf.comav.upv.es.

The authors declare that they have no competing financial interests that might have influenced the performance or presentation of the work described in this manuscript.

## Supplementary Tables

**Supplementary Table 1.** Accessions of domesticated and wild *Cucurbita* spp. used for transcriptomic and phylogenetic analyses. Number of reads used for assembly the transcriptomes, and number of genes and transcripts obtained are also shown.

| Code | Donor bank | Species | Subspecies (cultivar-group) | Country | Observations* | Number of reads | Number of genes | Number of transcripts |
| --- | --- | --- | --- | --- | --- | --- | --- | --- |
| BGV004370 | COMAV | *C. pepo* | *pepo* (Zucchini) | Spain |  | 1,892,580 | 18,446 | 18,902 |
| BGV005382 | COMAV | *C. pepo* | *ovifera* (Scallop) | Spain |  | 3,858,970 | 30,202 | 31,603 |
| PI 615111 | USDA | *C. pepo* | *ovifera* (Acorn) | USA |  | 7,305,712 | 43,585 | 47,003 |
| CATIE 18887 | CATIE | *C. pepo* | *pepo* (Pumpkin) | Mexico |  | 9,043,764 | 48,101 | 52,134 |
| CATIE 11368 | CATIE | *C. pepo* | *pepo* (Pumpkin) | Guatemala |  | 7,881,248 | 44,956 | 48,891 |
| PI 532354 | USDA | *C. pepo* | *fraterna* | Mexico |  | 14,574,156 | 54,631 | 63,051 |
| PI 614701 | USDA | *C. pepo* | *ozarkana* | USA | Reclassified as *fraterna* based on fruit traits | 11,131,674 | 41,507 | 45,626 |
| Nigerian Local | Seed company | *C. moschata* |  | Nigeria |  | 11,013,414 | 46,834 | 51,731 |
| PI 498429 | USDA | *C. moschata* |  | Colombia |  | 6,371,004 | 47,871 | 52,874 |
| PI 653064 | USDA | *C. moschata* |  | Nigeria |  | 8,793,992 | 40,308 | 43,531 |
| BGV004558 | COMAV | *C. maxima* |  | Argentina |  | 7,687,590 | 49,103 | 55,178 |
| VIR 3202 | VIR | *C. maxima* |  | Chile |  | 6,941,722 | 45,663 | 49,882 |
| UPV035142 | COMAV | *C. maxima* |  | Angola |  | 8,444,322 | 51,958 | 59,291 |
| PI 458653 | USDA | *C. maxima* | *andreana* | Argentina |  | 9,186,758 | 39,317 | 42,617 |
| PI 512115 | USDA | *C. argyrosperma* | *argyrosperma* | Guatemala |  | 8,431,320 | 44,179 | 50,119 |
| PI 451712 | USDA | *C. argyrosperma* | *argyrosperma* | USA |  | 17,537,688 | 56,350 | 69,160 |
| PI 438547 | USDA | *C. argyrosperma* |  | Belize |  | 13,654,318 | 50,299 | 58,029 |
| PI 202079 | USDA | *C. argyrosperma* | *argyrosperma* | Mexico |  | 10,173,060 | 43,676 | 48,547 |
| PI 512114 | USDA | *C. argyrosperma* | *argyrosperma* | Nicaragua |  | 8,312,838 | 37,028 | 40,099 |
| CATIE 16038 | CATIE | *C. ficifolia* |  | Guatemala |  | 11,304,986 | 40,213 | 43,269 |
| CATIE 16575 | CATIE | *C. ficifolia* |  | Guatemala |  | 11,415,514 | 54,537 | 60,549 |
| PI 432441 | USDA | *C. ecuadorensis* |  | Ecuador |  | 7,045,030 | 39,920 | 42,730 |
| PI 432443 | USDA | *C. ecuadorensis* |  | Ecuador |  | 5,165,038 | 43,184 | 46,323 |
| Grif 9446 | USDA | *C. ecuadorensis* |  | Ecuador |  | 11,424,844 | 49,330 | 54,999 |
| PI 532363 | USDA | *C. okeechobeensis* | *martinezii* | Mexico |  | 9,208,076 | 39,684 | 42,810 |
| PI 512105 | USDA | *C. okeechobeensis* | *martinezii* | Mexico |  | 5,670,572 | 42,321 | 45,035 |
| PI 512106 | USDA | *C. okeechobeensis* | *martinezii* | Mexico |  | 4,097,656 | 42,418 | 45,173 |
| PI 438542 | USDA | *C. lundelliana* |  | Belize |  | 10,515,350 | 45,096 | 49,266 |
| PI 532357 | USDA | *C. lundelliana* |  | Mexico |  | 6,957,802 | 40,933 | 44,365 |
| PI 636138 | USDA | *C. lundelliana* |  | Belize |  | 11,478,416 | 52,299 | 57,111 |
| PI 540898 | USDA | *C. lundelliana* |  | Honduras |  | 7,510,758 | 53,599 | 59,801 |
| PI 442197 | USDA | *C. foetidissima* |  | Mexico |  | 7,893,284 | 50,770 | 58,496 |
| PI 532350 | USDA | *C. foetidissima* |  | Mexico |  | 8,023,710 | 39,222 | 44,100 |
| PI 442201 | USDA | *C. foetidissima* |  | Mexico | Possible hybrid based on morphology | 12,257,620 | 54,205 | 65,217 |
| PI 532392 | USDA | *C. x scabridifolia* |  | Mexico | Hybrid *C. foetidissima* x *C. scabridifolia* | 9,427,190 | 42,885 | 48,184 |
| PI 653839 | USDA | *C. cordata* |  | Mexico |  | 5,055,088 | 38,784 | 41,705 |
| Grif 9445 | Seed company | *C. cordata* |  | Mexico |  | 5,756,634 | 33,530 | 36,153 |
| PI 442341 | USDA | *C. pedatifolia* |  | Mexico |  | 8,712,776 | 39,572 | 45,615 |
| PI 442290 | USDA | *C. pedatifolia* |  | Mexico |  | 27,822,108 | 67,366 | 92,522 |
| PI 540737 | USDA | *C. pedatifolia* |  | Mexico | Hybrid *C. pedatifolia* x *C. foetidissima* | 7,485,656 | 35,562 | 39,306 |

\* These accessions were morphologically characterized to confirm their taxonomic classification. Some of them were proved to be misclassified.

**Supplementary Table 2.** NGS library statistics. Numbers of raw reads, percentage of nucleotides over a 30 quality, coverage, % of reads filtered out during the cleaning process, % of reads without adaptor, % of chimeric reads, number of cleaned reads, coverage of cleaned reads, and percentage of nucleotides over a 30 quality in the clean reads.

| **Library** | **# Raw reads** | **Q30** | **Coverage** | **% Cleaned** | **% Category D** | **% Chimeric** | **# Filtered reads** | **Coverage clean** | **% Q30 clean** |
| --- | --- | --- | --- | --- | --- | --- | --- | --- | --- |
| Pair-end | 882,803,080 | 88.60 | 254 | 43.00 | -- | -- | 503,219,464 | 145 | 99.98 |
| 3 Kb | 186,878,960 | 91.32 | 54 | 49.60 | -- | 24.53 | 71,085,422 | 31 | 99.96 |
| 7 Kb | 159,602,336 | 90.74 | 46 | 73.67 | -- | 26.94 | 30,700,824 | 13 | 99.94 |
| 10 Kb | 149,980,526 | 89.62 | 65 | 48.40 | 28.59 | 3.55 | 53,381,028 | 23 | 99.986 |
| 20 Kb | 143,080,152 | 88.14 | 62 | 69.22 | 30.12 | 4.75 | 30,047,048 | 13 | 99.82 |

**Supplementary Table 3.**
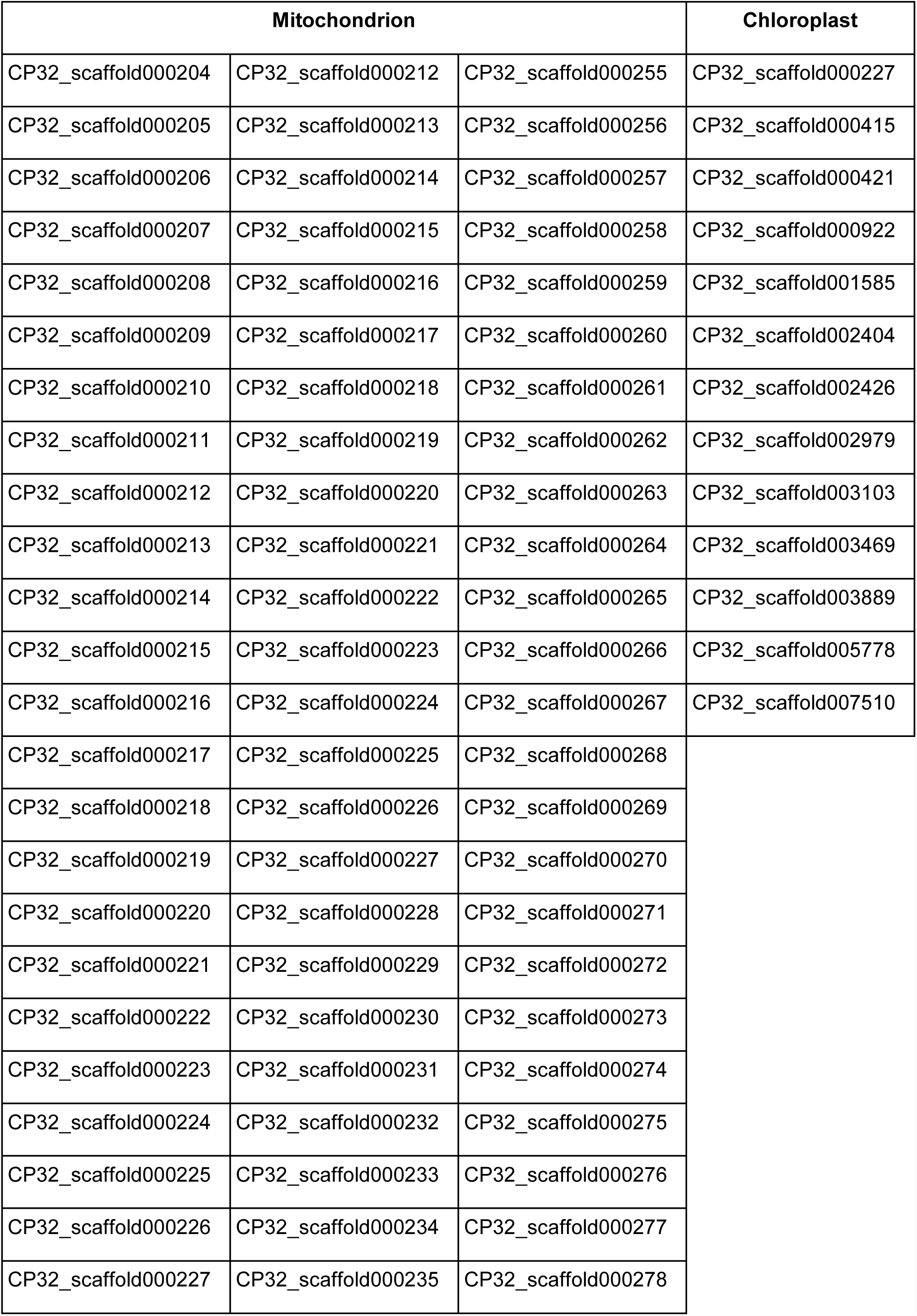
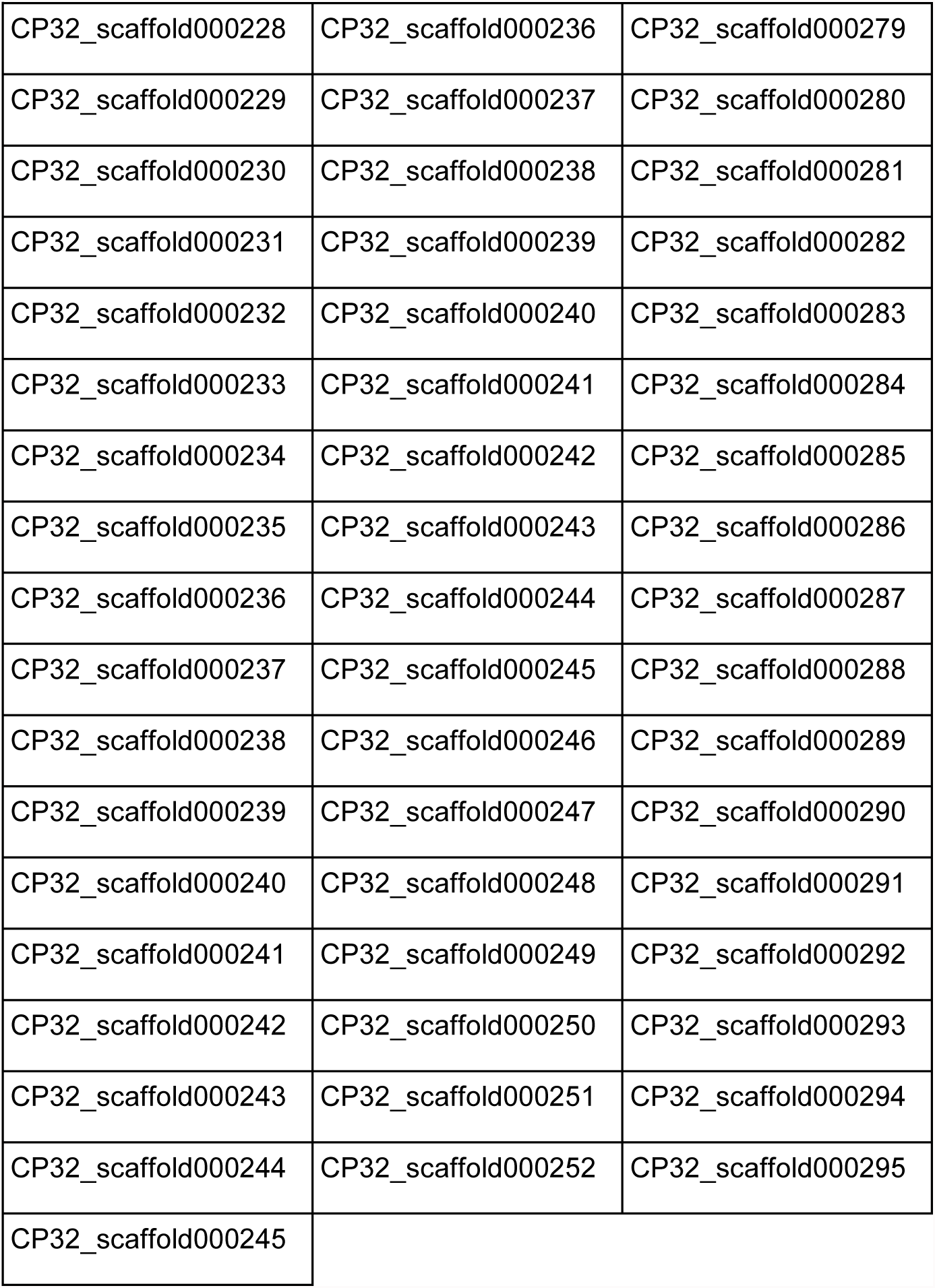
Scaffolds of genome assembly v.3.2. containing chloroplastic and mitochondrial regions. The pseudochromosomes were build out of the version 3.2 scaffolds.

| Mitochondrion |  |  | Chloroplast |
| --- | --- | --- | --- |
| CP32_scaffold000204 | CP32_scaffold000212 | CP32_scaffold000255 | CP32_scaffold000227 |
| CP32_scaffold000205 | CP32_scaffold000213 | CP32_scaffold000256 | CP32_scaffold000415 |
| CP32_scaffold000206 | CP32_scaffold000214 | CP32_scaffold000257 | CP32_scaffold000421 |
| CP32_scaffold000207 | CP32_scaffold000215 | CP32_scaffold000258 | CP32_scaffold000922 |
| CP32_scaffold000208 | CP32_scaffold000216 | CP32_scaffold000259 | CP32_scaffold001585 |
| CP32_scaffold000209 | CP32_scaffold000217 | CP32_scaffold000260 | CP32_scaffold002404 |
| CP32_scaffold000210 | CP32_scaffold000218 | CP32_scaffold000261 | CP32_scaffold002426 |
| CP32_scaffold000211 | CP32_scaffold000219 | CP32_scaffold000262 | CP32_scaffold002979 |
| CP32_scaffold000212 | CP32_scaffold000220 | CP32_scaffold000263 | CP32_scaffold003103 |
| CP32_scaffold000213 | CP32_scaffold000221 | CP32_scaffold000264 | CP32_scaffold003469 |
| CP32_scaffold000214 | CP32_scaffold000222 | CP32_scaffold000265 | CP32_scaffold003889 |
| CP32_scaffold000215 | CP32_scaffold000223 | CP32_scaffold000266 | CP32_scaffold005778 |
| CP32_scaffold000216 | CP32_scaffold000224 | CP32_scaffold000267 | CP32_scaffold007510 |
| CP32_scaffold000217 | CP32_scaffold000225 | CP32_scaffold000268 |  |
| CP32_scaffold000218 | CP32_scaffold000226 | CP32_scaffold000269 |  |
| CP32_scaffold000219 | CP32_scaffold000227 | CP32_scaffold000270 |  |
| CP32_scaffold000220 | CP32_scaffold000228 | CP32_scaffold000271 |  |
| CP32_scaffold000221 | CP32_scaffold000229 | CP32_scaffold000272 |  |
| CP32_scaffold000222 | CP32_scaffold000230 | CP32_scaffold000273 |  |
| CP32_scaffold000223 | CP32_scaffold000231 | CP32_scaffold000274 |  |
| CP32_scaffold000224 | CP32_scaffold000232 | CP32_scaffold000275 |  |
| CP32_scaffold000225 | CP32_scaffold000233 | CP32_scaffold000276 |  |
| CP32_scaffold000226 | CP32_scaffold000234 | CP32_scaffold000277 |  |
| CP32_scaffold000227 | CP32_scaffold000235 | CP32_scaffold000278 |  |

|  |  |  |
| --- | --- | --- |
| CP32_scaffold000228 | CP32_scaffold000236 | CP32_scaffold000279 |
| CP32_scaffold000229 | CP32_scaffold000237 | CP32_scaffold000280 |
| CP32_scaffold000230 | CP32_scaffold000238 | CP32_scaffold000281 |
| CP32_scaffold000231 | CP32_scaffold000239 | CP32_scaffold000282 |
| CP32_scaffold000232 | CP32_scaffold000240 | CP32_scaffold000283 |
| CP32_scaffold000233 | CP32_scaffold000241 | CP32_scaffold000284 |
| CP32_scaffold000234 | CP32_scaffold000242 | CP32_scaffold000285 |
| CP32_scaffold000235 | CP32_scaffold000243 | CP32_scaffold000286 |
| CP32_scaffold000236 | CP32_scaffold000244 | CP32_scaffold000287 |
| CP32_scaffold000237 | CP32_scaffold000245 | CP32_scaffold000288 |
| CP32_scaffold000238 | CP32_scaffold000246 | CP32_scaffold000289 |
| CP32_scaffold000239 | CP32_scaffold000247 | CP32_scaffold000290 |
| CP32_scaffold000240 | CP32_scaffold000248 | CP32_scaffold000291 |
| CP32_scaffold000241 | CP32_scaffold000249 | CP32_scaffold000292 |
| CP32_scaffold000242 | CP32_scaffold000250 | CP32_scaffold000293 |
| CP32_scaffold000243 | CP32_scaffold000251 | CP32_scaffold000294 |
| CP32_scaffold000244 | CP32_scaffold000252 | CP32_scaffold000295 |
| CP32_scaffold000245 |  |  |

**Supplementary Table 4.** Genome v.4.1 pseudochromosomes configuration. The order, orientation and size of genome v. 3.2 scaffolds grouped in each pseudochromosomes is shown. Equivalence of pseudochromosomes and linkage groups of Montero-Pau et al. (2016) genetic map is also shown.

| **Genome v.4.1**<br>**pseudochromosome** | **Genetic map linkage**<br>**group in Montero-Pau et**<br>**al. 2016** | **Scaffold**<br>**order** | **Scaffold name** | **Scaffold**<br>**size** | **Scaffold**<br>**orientation** |
| --- | --- | --- | --- | --- | --- |
| CP4.1LG01 | LG01 | 1 | CP32_scaffold000010 | 3,883,160 | reverse |
|  |  | 2 | CP32_scaffold000175 | 233,039 | reverse |
|  |  | 3 | CP32_scaffold000046 | 1,602,464 | reverse |
|  |  | 4 | CP32_scaffold000051 | 1,450,516 | reverse |
|  |  | 5 | CP32_scaffold000040 | 1,852,384 | reverse |
|  |  | 6 | CP32_scaffold000144 | 331,714 | forward |
|  |  | 7 | CP32_scaffold000084 | 741,359 | forward |
|  |  | 8 | CP32_scaffold000063 | 1,087,272 | forward |
|  |  | 9 | CP32_scaffold000111 | 460,681 | forward |
|  |  | 10 | CP32_scaffold000233 | 109,860 | reverse |
|  |  | 11 | CP32_scaffold000078 | 832,487 | forward |
|  |  | 12 | CP32_scaffold000181 | 222,517 | reverse |
|  |  | 13 | CP32_scaffold000059 | 1,137,078 | reverse |
|  |  | 14 | CP32_scaffold000079 | 790,494 | forward |
|  |  | 15 | CP32_scaffold000087 | 658,871 | reverse |
|  |  | 16 | CP32_scaffold000105 | 507,762 | forward |
|  |  | 17 | CP32_scaffold000032 | 2,241,395 | forward |
|  |  | 18 | CP32_scaffold000024 | 2,525,860 | reverse |
|  |  | 19 | CP32_scaffold000091 | 633,856 | forward |
| CP4.1LG02 | LG02 | 1 | CP32_scaffold000001 | 6,123,784 | reverse |
|  |  | 2 | CP32_scaffold000128 | 403,365 | reverse |
|  |  | 3 | CP32_scaffold000099 | 554,539 | forward |
|  |  | 4 | CP32_scaffold000166 | 257,955 | forward |
|  |  | 5 | CP32_scaffold000090 | 642,088 | undefined |
|  |  | 6 | CP32_scaffold000076 | 841,349 | undefined |
|  |  | 7 | CP32_scaffold000055 | 1,317,493 | reverse |
|  |  | 8 | CP32_scaffold000162 | 279,451 | reverse |
|  |  | 9 | CP32_scaffold000208 | 154,271 | forward |
|  |  | 10 | CP32_scaffold000374 | 40,295 | undefined |
|  |  | 11 | CP32_scaffold000140 | 338,305 | forward |
|  |  | 12 | CP32_scaffold000062 | 1,102,807 | reverse |
|  |  | 13 | CP32_scaffold002675 | 2,094 | undefined |
|  |  | 14 | CP32_scaffold000169 | 245,895 | reverse |
|  |  | 15 | CP32_scaffold000041 | 1,808,812 | reverse |
|  |  | 16 | CP32_scaffold000168 | 248,911 | forward |
| CP4.1LG03 | LG03 | 1 | CP32_scaffold000038 | 1,877,481 | reverse |
|  |  | 2 | CP32_scaffold000122 | 422,066 | reverse |
|  |  | 3 | CP32_scaffold000195 | 188,662 | forward |
|  |  | 4 | CP32_scaffold000019 | 3,089,961 | reverse |
|  |  | 5 | CP32_scaffold000117 | 430,983 | reverse |
|  |  | 6 | CP32_scaffold000210 | 151,564 | forward |
|  |  | 7 | CP32_scaffold000163 | 278,935 | forward |
|  |  | 8 | CP32_scaffold000177 | 230,436 | undefined |
|  |  | 9 | CP32_scaffold000118 | 428,730 | forward |
|  |  | 10 | CP32_scaffold000187 | 210,711 | forward |
|  |  | 11 | CP32_scaffold000006 | 4,754,185 | reverse |
|  |  | 12 | CP32_scaffold000044 | 1,697,700 | reverse |
| CP4.1LG04 | LG18 + LG20 | 1 | CP32_scaffold000027 | 2,403,537 | reverse |
|  |  | 2 | CP32_scaffold000207 | 155,442 | forward |
|  |  | 3 | CP32_scaffold000002 | 5,399,389 | reverse |
|  |  | 4 | CP32_scaffold000025 | 2,522,528 | reverse |
|  |  | 5 | CP32_scaffold000033 | 2,224,244 | forward |
| CP4.1LG05 | LG04 | 1 | CP32_scaffold000066 | 995,157 | reverse |
|  |  | 2 | CP32_scaffold000180 | 223,211 | reverse |
|  |  | 3 | CP32_scaffold000143 | 334,696 | forward |
|  |  | 4 | CP32_scaffold000086 | 693,282 | reverse |
|  |  | 5 | CP32_scaffold000159 | 289,565 | forward |
|  |  | 6 | CP32_scaffold000022 | 2,747,443 | forward |
|  |  | 7 | CP32_scaffold000021 | 2,780,951 | forward |
|  |  | 8 | CP32_scaffold000020 | 2,794,373 | forward |
| CP4.1LG06 | LG07 | 1 | CP32_scaffold000035 | 2,140,805 | forward |
|  |  | 2 | CP32_scaffold000083 | 758,721 | reverse |
|  |  | 3 | CP32_scaffold000042 | 1,749,822 | forward |
|  |  | 4 | CP32_scaffold000043 | 1,732,427 | reverse |
|  |  | 5 | CP32_scaffold000082 | 760,949 | reverse |
|  |  | 6 | CP32_scaffold000185 | 216,892 | forward |
|  |  | 7 | CP32_scaffold013127 | 333 | forward |
|  |  | 8 | CP32_scaffold000158 | 289,633 | reverse |
|  |  | 9 | CP32_scaffold000036 | 2,105,938 | reverse |
|  |  | 10 | CP32_scaffold000106 | 503,388 | forward |
|  |  | 11 | CP32_scaffold000127 | 408,837 | forward |
| CP4.1LG07 | LG15 | 1 | CP32_scaffold000060 | 1,133,474 | forward |
|  |  | 2 | CP32_scaffold000093 | 615,043 | reverse |
|  |  | 3 | CP32_scaffold000056 | 1,254,317 | forward |
|  |  | 4 | CP32_scaffold000097 | 558,874 | reverse |
|  |  | 5 | CP32_scaffold000069 | 945,996 | reverse |
|  |  | 6 | CP32_scaffold000135 | 366,381 | reverse |
|  |  | 7 | CP32_scaffold000119 | 428,542 | reverse |
|  |  | 8 | CP32_scaffold000133 | 378,800 | undefined |
|  |  | 9 | CP32_scaffold000023 | 2,545,012 | forward |
|  |  | 10 | CP32_scaffold000138 | 352,159 | reverse |
|  |  | 11 | CP32_scaffold000214 | 135,984 | forward |
|  |  | 12 | CP32_scaffold000172 | 241,243 | reverse |
|  |  | 13 | CP32_scaffold000092 | 623,336 | forward |
|  |  | 14 | CP32_scaffold000098 | 555,395 | reverse |
| CP4.1LG08 | LG06 | 1 | CP32_scaffold000003 | 4,875,806 | reverse |
|  |  | 2 | CP32_scaffold000089 | 642,205 | forward |
|  |  | 3 | CP32_scaffold000008 | 3,995,464 | forward |
|  |  | 4 | CP32_scaffold000101 | 542,828 | forward |
| CP4.1LG09 | LG08 | 1 | CP32_scaffold000014 | 3,564,952 | forward |
|  |  | 2 | CP32_scaffold000251 | 101,034 | forward |
|  |  | 3 | CP32_scaffold000061 | 1,116,870 | reverse |
|  |  | 4 | CP32_scaffold000132 | 379,208 | reverse |
|  |  | 5 | CP32_scaffold000245 | 101,953 | forward |
|  |  | 6 | CP32_scaffold000247 | 101,524 | reverse |
|  |  | 7 | CP32_scaffold000145 | 331,494 | undefined |
|  |  | 8 | CP32_scaffold000075 | 863,406 | forward |
|  |  | 9 | CP32_scaffold000072 | 896,126 | forward |
|  |  | 10 | CP32_scaffold000026 | 2,454,755 | forward |
| CP4.1LG10 | LG10 | 1 | CP32_scaffold000009 | 3,958,430 | reverse |
|  |  | 2 | CP32_scaffold000029 | 2,358,218 | reverse |
|  |  | 3 | CP32_scaffold000068 | 950,888 | reverse |
|  |  | 4 | CP32_scaffold000125 | 410,804 | forward |
|  |  | 5 | CP32_scaffold000160 | 285,501 | forward |
|  |  | 6 | CP32_scaffold000057 | 1,171,448 | reverse |
|  |  | 7 | CP32_scaffold000171 | 242,204 | reverse |
|  |  | 8 | CP32_scaffold000114 | 450,599 | reverse |
| CP4.1LG11 | LG13 | 1 | CP32_scaffold000107 | 498,007 | reverse |
|  |  | 2 | CP32_scaffold000017 | 3,213,514 | forward |
|  |  | 3 | CP32_scaffold000108 | 488,883 | forward |
|  |  | 4 | CP32_scaffold000124 | 411,803 | reverse |
|  |  | 5 | CP32_scaffold000141 | 338,006 | forward |
|  |  | 6 | CP32_scaffold000088 | 647,301 | reverse |
|  |  | 7 | CP32_scaffold000113 | 452,942 | reverse |
|  |  | 8 | CP32_scaffold000049 | 1,492,573 | forward |
|  |  | 9 | CP32_scaffold000053 | 1,373,939 | forward |
|  |  | 10 | CP32_scaffold000094 | 602,087 | forward |
|  |  | 11 | CP32_scaffold000153 | 304,914 | forward |
| CP4.1LG12 | LG05 | 1 | CP32_scaffold000012 | 3,815,302 | forward |
|  |  | 2 | CP32_scaffold000121 | 422,947 | reverse |
|  |  | 3 | CP32_scaffold000071 | 899,720 | forward |
|  |  | 4 | CP32_scaffold000018 | 3,164,343 | reverse |
|  |  | 5 | CP32_scaffold000048 | 1,517,882 | reverse |
| CP4.1LG13 | LG16 | 1 | CP32_scaffold000085 | 695,292 | forward |
|  |  | 2 | CP32_scaffold000034 | 2,168,721 | forward |
|  |  | 3 | CP32_scaffold000129 | 385,840 | forward |
|  |  | 4 | CP32_scaffold000225 | 119,458 | reverse |
|  |  | 5 | CP32_scaffold000080 | 770,750 | forward |
|  |  | 6 | CP32_scaffold000028 | 2,396,146 | reverse |
|  |  | 7 | CP32_scaffold000030 | 2,292,851 | reverse |
|  |  | 8 | CP32_scaffold000104 | 518,031 | forward |
| CP4.1LG14 | LG19 | 1 | CP32_scaffold000005 | 4,849,021 | reverse |
|  |  | 2 | CP32_scaffold000147 | 327,288 | undefined |
|  |  | 3 | CP32_scaffold000182 | 222,078 | undefined |
|  |  | 4 | CP32_scaffold000050 | 1,454,989 | forward |
|  |  | 5 | CP32_scaffold000037 | 2,098,557 | forward |
| CP4.1LG15 | LG11 | 1 | CP32_scaffold000015 | 3,441,236 | forward |
|  |  | 2 | CP32_scaffold000131 | 379,828 | forward |
|  |  | 3 | CP32_scaffold000047 | 1,558,130 | forward |
|  |  | 4 | CP32_scaffold000016 | 3,434,250 | forward |
| CP4.1LG16 | LG17 | 1 | CP32_scaffold000067 | 991,265 | reverse |
|  |  | 2 | CP32_scaffold000200 | 168,863 | forward |
|  |  | 3 | CP32_scaffold000164 | 278,388 | undefined |
|  |  | 4 | CP32_scaffold000256 | 97,354 | undefined |
|  |  | 5 | CP32_scaffold000070 | 928,547 | forward |
|  |  | 6 | CP32_scaffold000206 | 156,815 | undefined |
|  |  | 7 | CP32_scaffold000152 | 310,330 | forward |
|  |  | 8 | CP32_scaffold000100 | 550,348 | forward |
|  |  | 9 | CP32_scaffold000142 | 337,580 | forward |
|  |  | 10 | CP32_scaffold000004 | 4,863,444 | forward |
| CP4.1LG17 | LG12 | 1 | CP32_scaffold000103 | 531,021 | reverse |
|  |  | 2 | CP32_scaffold000065 | 997,141 | forward |
|  |  | 3 | CP32_scaffold000074 | 869,430 | reverse |
|  |  | 4 | CP32_scaffold000096 | 589,541 | forward |
|  |  | 5 | CP32_scaffold000120 | 423,438 | reverse |
|  |  | 6 | CP32_scaffold000186 | 215,517 | forward |
|  |  | 7 | CP32_scaffold000058 | 1,167,845 | forward |
|  |  | 8 | CP32_scaffold000031 | 2,250,994 | reverse |
|  |  | 9 | CP32_scaffold000045 | 1,627,577 | forward |
| CP4.1LG18 | LG14 | 1 | CP32_scaffold000073 | 884,058 | forward |
|  |  | 2 | CP32_scaffold000095 | 591,145 | forward |
|  |  | 3 | CP32_scaffold000116 | 432,341 | forward |
|  |  | 4 | CP32_scaffold000157 | 293,129 | forward |
|  |  | 5 | CP32_scaffold000112 | 453,344 | forward |
|  |  | 6 | CP32_scaffold000039 | 1,853,076 | forward |
|  |  | 7 | CP32_scaffold000011 | 3,820,361 | forward |
| CP4.1LG19 | LG09 | 1 | CP32_scaffold000054 | 1,345,344 | forward |
|  |  | 2 | CP32_scaffold000064 | 1,081,996 | forward |
|  |  | 3 | CP32_scaffold000161 | 281,296 | forward |
|  |  | 4 | CP32_scaffold000197 | 183,073 | reverse |
|  |  | 5 | CP32_scaffold000146 | 330,575 | forward |
|  |  | 6 | CP32_scaffold000077 | 836,762 | reverse |
|  |  | 7 | CP32_scaffold000130 | 382,378 | reverse |
|  |  | 8 | CP32_scaffold000013 | 3,798,258 | reverse |
| CP4.1LG20 | LG21 | 1 | CP32_scaffold000007 | 4,346,540 | forward |
|  |  | 2 | CP32_scaffold000148 | 327,036 | reverse |
|  |  | 3 | CP32_scaffold000115 | 449,779 | forward |
|  |  | 4 | CP32_scaffold000110 | 469,294 | forward |
|  |  | 5 | CP32_scaffold000151 | 313,452 | forward |
|  |  | 6 | CP32_scaffold000081 | 769,043 | reverse |
|  |  | 7 | CP32_scaffold000052 | 1,439,660 | reverse |

**Supplementary Table 5.** Summary of repetitive elements found in *Cucurbita pepo, Cucumis melo, Cucumis sativus* and *Citrullus lanatus.* All results are expressed in bp.

| Size (bp) | *Cucurbita*<br>*pepo* | *Cucumis*<br>*melo* | *Citrullus*<br>*lanatus* | *Cucumis*<br>*sativus*<br>Chinese long | *Cucumis*<br>*sativus*<br>PI183967 |
| --- | --- | --- | --- | --- | --- |
| Genome size | 289504453 | 406928820 | 355247419 | 197271687 | 204803225 |
| Genome size without Ns | 247816929 | 336097957 | 321405453 | 193700889 | 200988521 |
| Repetitive | 93650597 | 190225685 | 186381889 | 55566601 | 63676019 |
| Repetitive (no overlapping) | 85581680 | 179982107 | 173126881 | 52923711 | 60250642 |
| over_same | 4361793 | 5007047 | 5992548 | 1199958 | 1697636 |
| over_diff | 3707124 | 5236531 | 7262460 | 1442932 | 1727741 |
| DNA |  | 193677 | 283426 | 54557 |  |
| DNA/CMC-EnSpm | 88523 | 11363266 | 2589519 | 1551581 | 1955210 |
| DNA/Crypton-C |  |  | 7193 |  |  |
| DNA/En-Spm |  |  | 139908 |  |  |
| DNA/Ginger |  |  | 121896 | 134666 |  |
| DNA/IS3EU |  |  |  |  | 82374 |
| DNA/Kolobok-T2 | 27949 |  |  |  | 58226 |
| DNA/MULE-MuDR | 145882 | 4697101 | 1633726 | 1195740 | 1003127 |
| DNA/Maverick |  |  | 192173 |  |  |
| DNA/MuDR |  | 929167 | 122428 | 136004 | 148317 |
| DNA/MuLE-MuDR |  | 6548932 | 4029767 | 890154 | 996088 |
| DNA/PIF-Harbinger | 12213 | 3090436 | 1290961 | 550468 | 343060 |
| DNA/TcMar-Mariner |  |  | 3360 |  |  |
| DNA/Zisupton |  |  |  | 25310 |  |
| DNA/hAT-Ac | 1246686 | 728666 | 1709109 | 264199 | 479104 |
| DNA/hAT-Charlie |  | 148193 | 38283 | 60790 |  |
| DNA/hAT-Tag1 | 159459 | 383200 | 478025 | 345958 | 361065 |
| DNA/hAT-Tip100 |  | 379594 | 83286 | 367630 | 83315 |
| DNA/hAT-hATm |  |  | 2825369 |  |  |
| LINE/CR1 |  |  |  | 36758 | 21279 |
| LINE/CRE | 169102 |  | 31891 | 20041 |  |
| LINE/I-Jockey |  | 301 |  |  |  |
| LINE/L1 | 2207438 | 5388043 | 5944566 | 2957971 | 3237632 |
| LINE/L1-DRE |  |  | 217246 |  | 86388 |
| LINE/L1-Tx1 | 194920 | 201316 | 224728 | 23383 |  |
| LINE/L2 |  | 24417 | 950717 | 70984 |  |
| LINE/RTE-BovB | 143 |  | 185123 |  | 104075 |
| LINE/Tad1 | 68219 |  |  |  | 117729 |
| LTR | 65446 | 200156 | 263981 | 334749 |  |
| LTR/Cassandra | 1310467 | 387478 | 346831 | 236843 | 41046 |
| LTR/Caulimovirus | 719038 | 1100094 | 225529 | 122829 | 55250 |
| LTR/Copia | 6667202 | 39238863 | 33496488 | 9503582 | 10759234 |
| LTR/DIRS | 150502 |  |  |  |  |
| LTR/ERV1 |  | 189116 |  |  | 12581 |
| LTR/ERVK | 30609 |  |  |  |  |
| LTR/ERVL |  |  | 136788 |  |  |
| LTR/Gypsy | 8160792 | 43716718 | 31453573 | 5844651 | 7988269 |
| LTR/Pao |  | 12326 |  |  | 32675 |
| RC/Helitron | 181502 | 632496 | 114736 | 71766 | 48395 |
| Retroposon |  |  |  | 73613 |  |
| SINE/Alu |  |  |  |  | 48743 |
| SINE/B2 | 11862 |  | 11186 |  |  |
| SINE/ID | 175670 |  |  |  |  |
| SINE/tRNA | 200088 | 333231 | 686539 | 255170 | 246352 |
| SINE/tRNA-R2 | 23160 |  |  |  |  |
| SINE/tRNA-RTE | 76347 |  |  |  |  |
| SINE? | 1376290 | 2458 | 20168 |  | 41669 |
| Satellite | 399460 | 232131 |  | 13933 | 6340 |
| Satellite/Y-chromosome | 31503 |  |  |  |  |
| Satellite/centr |  |  |  |  | 63047 |
| Simple repeats | 8092785 | 5360259 | 5874319 | 3676307 | 4190723 |
| Low complexity regions | 1229916 | 1395906 | 1646367 | 1175901 | 1160849 |
| rRNA | 468015 | 176725 | 41472 | 50262 | 109384 |
| snRNA | 29907 | 7471 | 6485 | 6750 | 4777 |
| Unknown/undefined | 59929502 | 63163948 | 88954727 | 25514051 | 29789696 |

**Supplementary Table 6.** Gene family (orthogroups and paralogs in OrthoMCL) identification.

| **Species** | **# proteins** | **# assigned to**<br>**a gene family**<br>**(several**<br>**species)** | **% assigned** | **# assigned to**<br>**a gene family**<br>**(species**<br>**exclusive)** | **% assigned to**<br>**a gene family**<br>**(species**<br>**exclusive)** | **% of**<br>**proteins**<br>**assigned** |
| --- | --- | --- | --- | --- | --- | --- |
| *Cucurbita pepo* | 27,870 | 25,433 | 91.23 | 291 | 1.04 | 95.26 |
| *Citrullus lanatus* | 23,440 | 18,798 | 80.20 | 2,601 | 11.10 | 94.16 |
| *Cucumis melo* | 27,427 | 19,974 | 72.83 | 3,570 | 13.02 | 88.22 |
| *Cucumis sativus*<br>Chinese Long | 23,248 | 19,111 | 82.20 | 394 | 1.69 | 86.94 |
| *Cucumis sativus*<br>PI183967 | 22,790 | 19,360 | 84.95 | 370 | 1.62 | 89.48 |

Supplementary Table 7. List of genes that are single copy in *Cucurbita pepo.* Predicted function is also shown.

Supplementary Table 8. List of genes that are duplicated in *Cucurbita pepo.* Predicted function is also shown.

Supplementary Table 9. List of genes that are duplicated in *Cucurbita pepo* but not in *Cucumis melo, Cucumis sativus* or *Citrullus lanatus.* Predicted function is also shown.

Supplementary Table 10. GO term enrichment tests. Results are shown for single copy genes in *Cucurbita pepo,* duplicated genes, and genes that are exclusively duplicated in *C. pepo* when compared with other cucurbit genomes.

## Supplementary Images

**Supplementary Figure 1.**
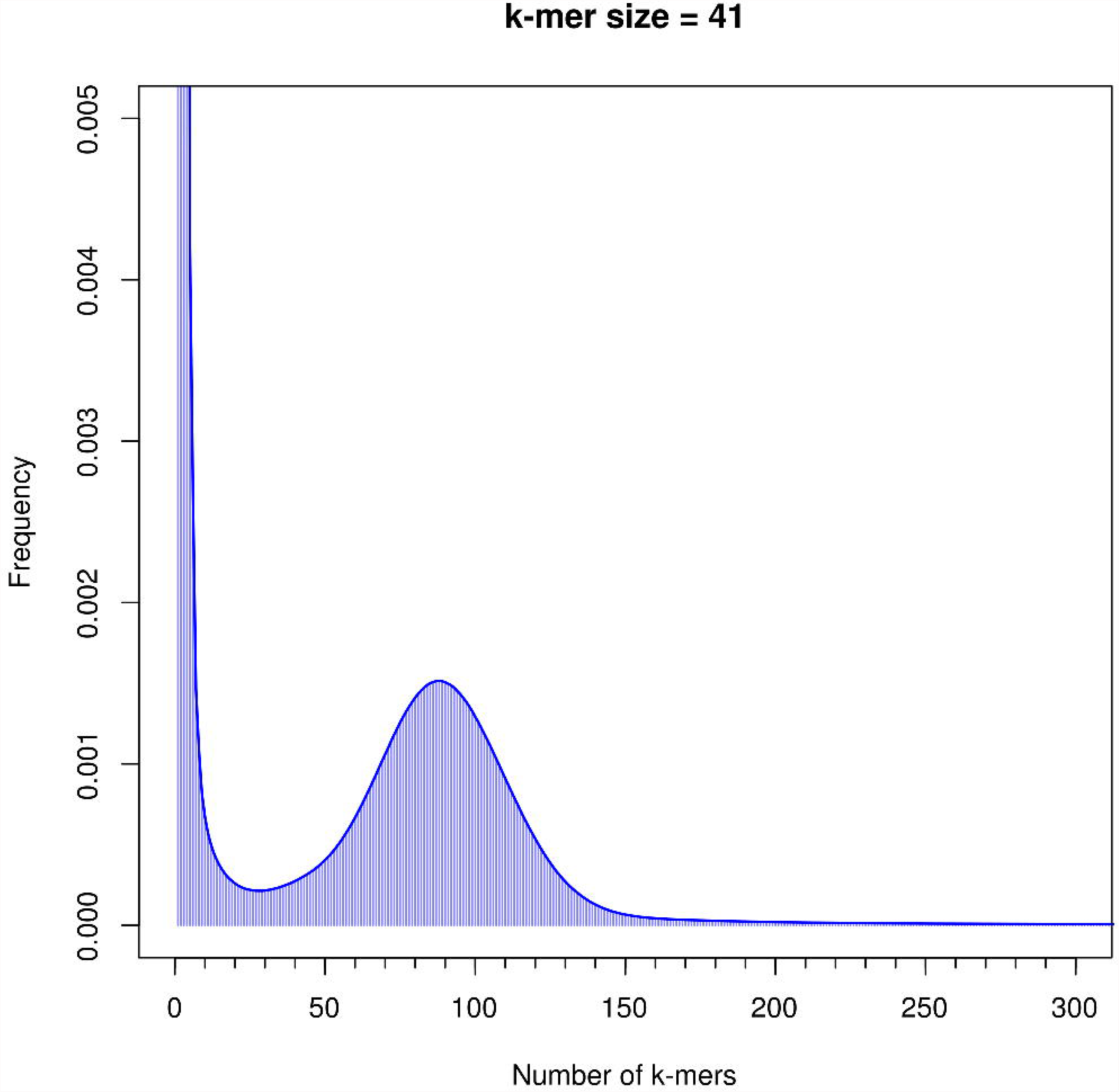
Distribution of sequences of k-mer size 41 for different levels of coverage

**Supplementary Figure 2.**
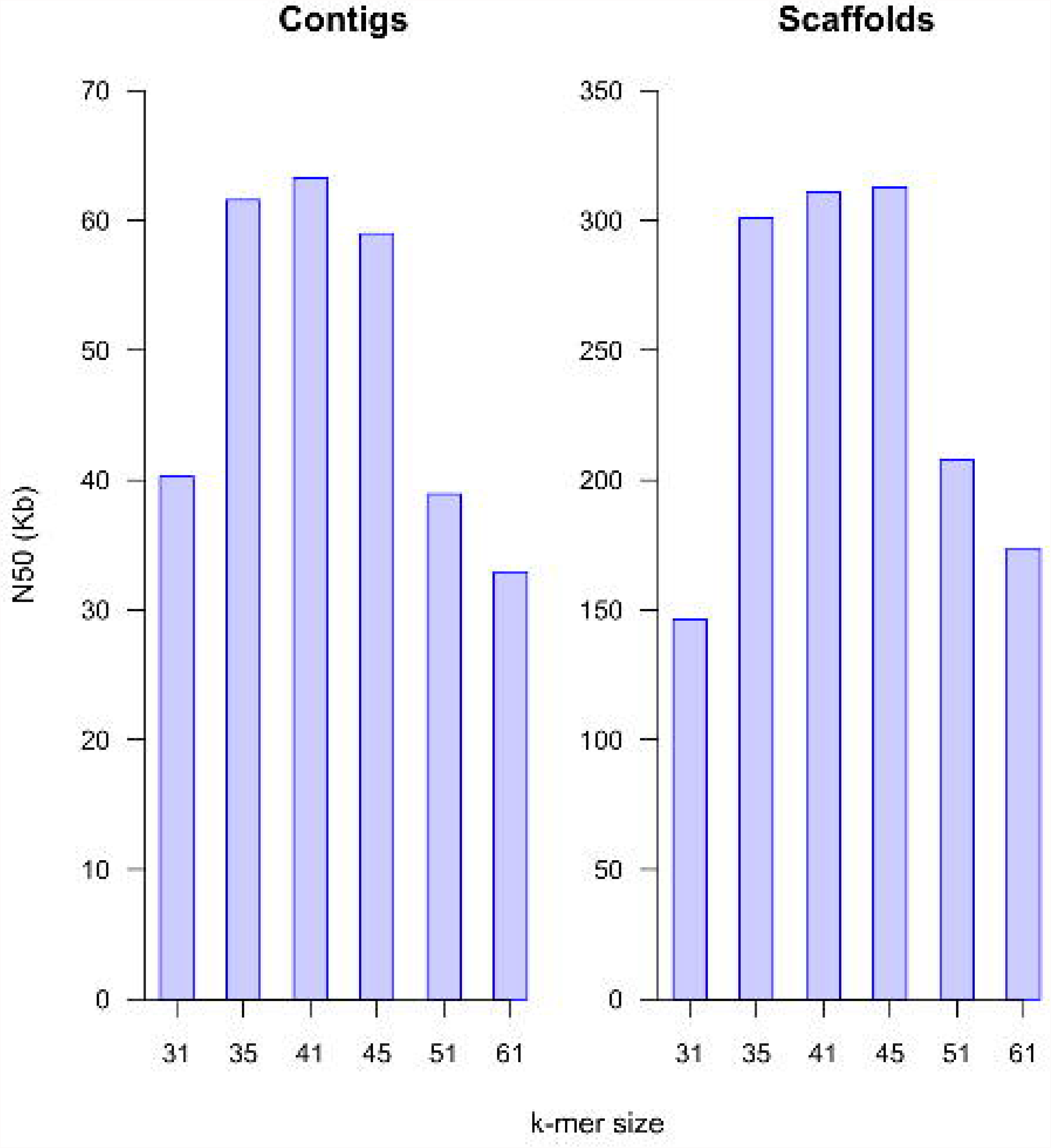
Distribution of N50 for contigs (A) and scaffolds (B) for different k-mer size values

**Supplementary Figure 3.**
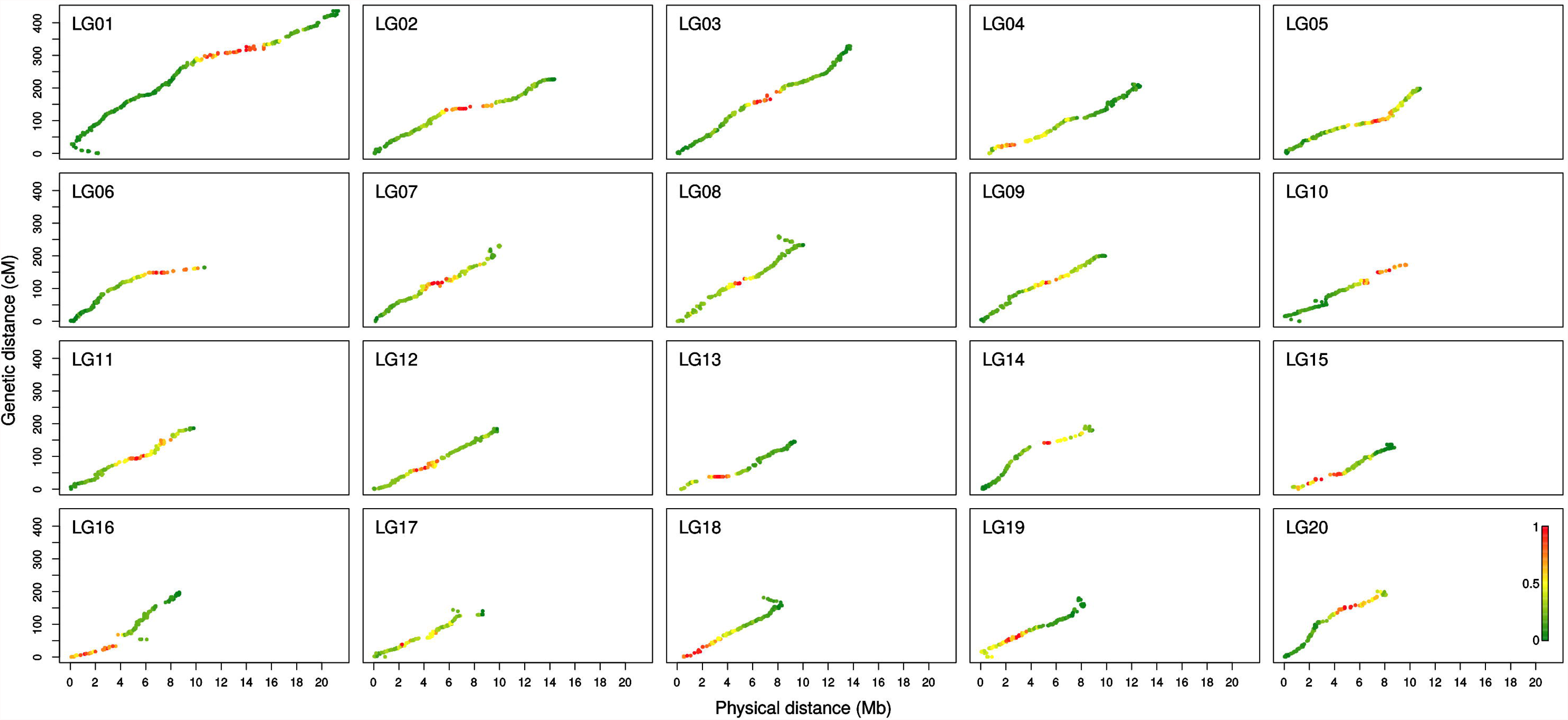
Correlation between genetic and physical distances for each pseudochromosome. Color scale represents fraction of repetitive DNA.

**Supplementary Figure 4.**
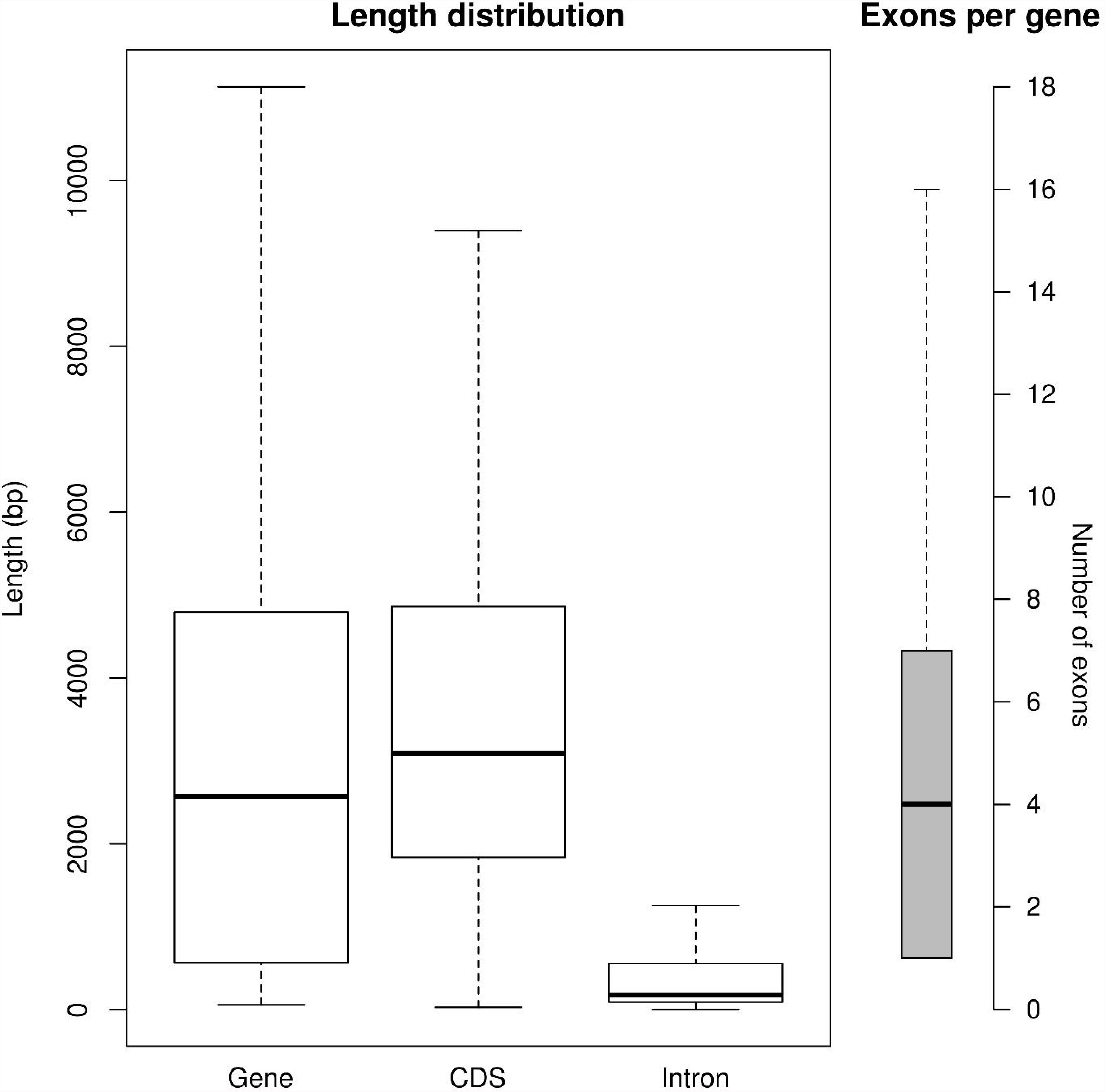
Summary of the structural annotation of *C. pepo* genome.

**Supplementary Figure 5.**
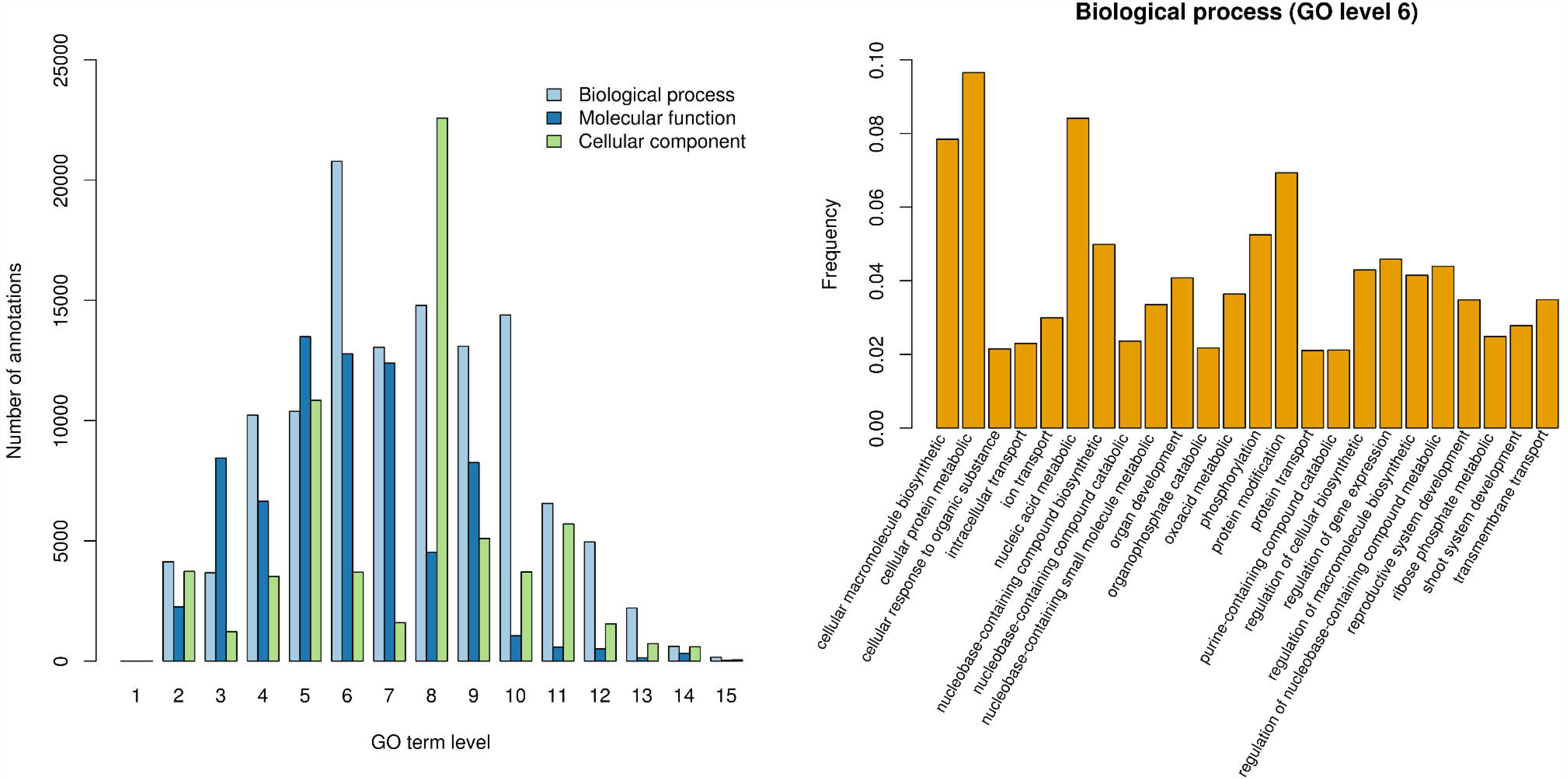
Transcriptome GO annotation statistics A) by levels and B) at level 6.

**Supplementary Figure 6.**
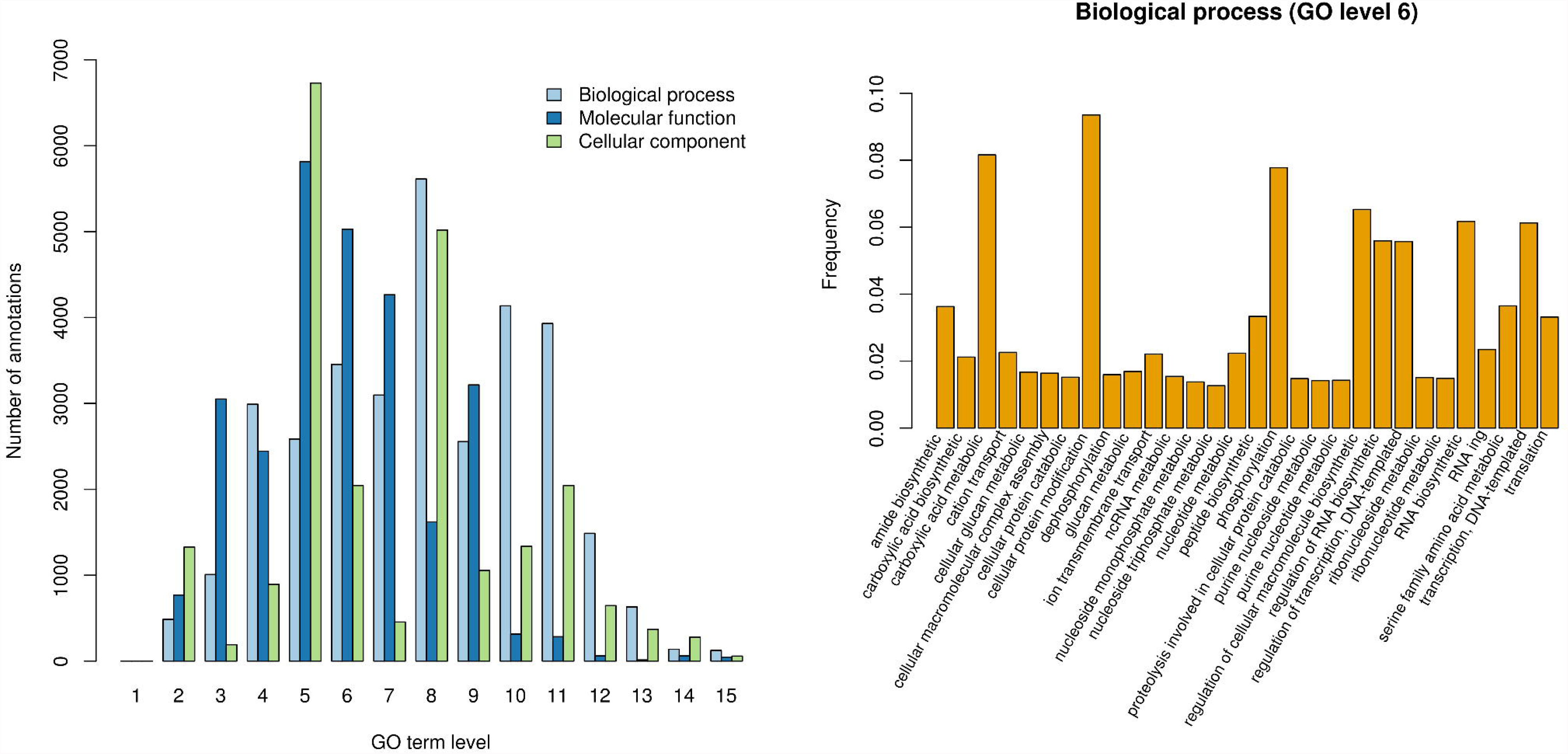
Genome GO annotation statistics A) by levels and B) at level 6.

**Supplementary Figure 7.**
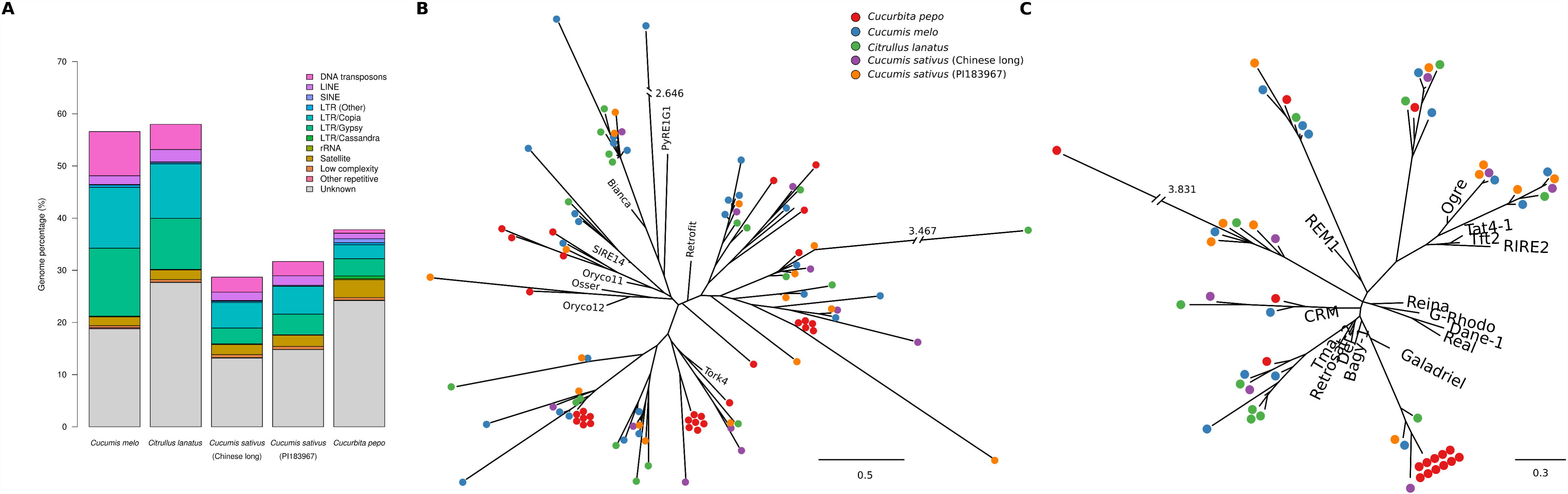
Repetitive elements. Fraction of genome covered by different types of repetitive elements in *C. pepo and* four *Cucurbita* genomes (A). Maximum likelihood phylogenetic trees of *C. pepo* elements of *Copia* (B) and *Gypsy* (C) LTR superfamilies based on a fragment of the reverse transcriptase.

**Supplementary Figure 8.**
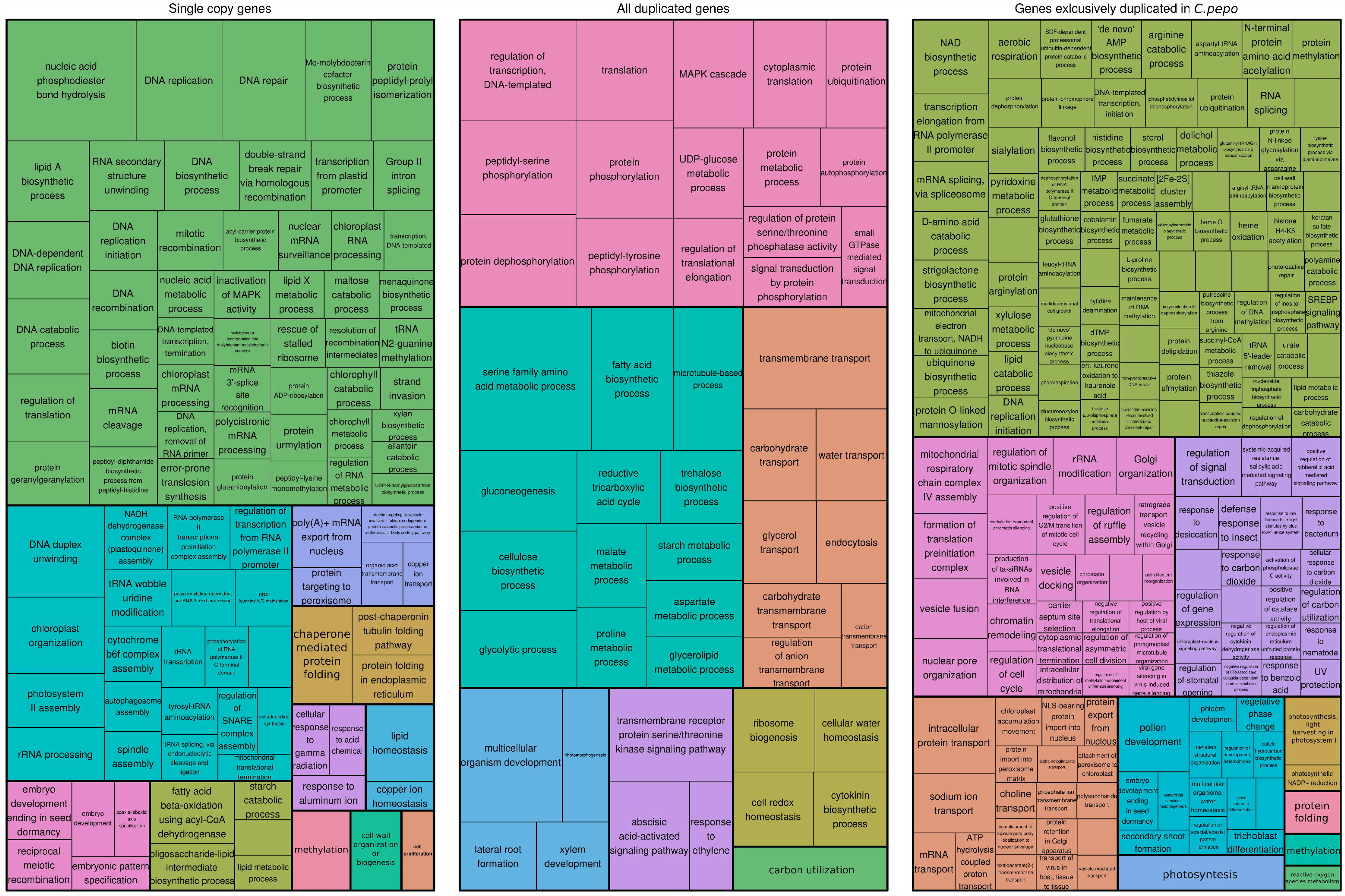
Results of GO enrichment test. Treemaps for the results of the GO enrichment tests on single copy genes in *Cucurbita pepo,* all duplicated genes in *C. pepo* and genes that are duplicated in *C. pepo* but not in other cucurbit genomes. Area of rectangles represent minus logarithm of enrichment test FDR.

## Supplementary data

Supplementary data 1. *Cucurbita pepo* genome assembly 4.1. Fasta file.

Supplementary data 2. *Cucurbita pepo* genome annotation 4.1. GFF file.

Supplementary data 3. *Cucurbita pepo* GO term annotation. Results from Blast2GO.

